# PAK1 and PAK2 in cell metabolism regulation

**DOI:** 10.1101/2021.05.18.444625

**Authors:** Tereza Kořánová, Lukáš Dvořáček, Dana Grebeňová, Pavla Röselová, Adam Obr, Kateřina Kuželová

## Abstract

P21-activated kinases (PAKs) regulate processes associated with cytoskeletal rearrangements, such as cell division, adhesion, and migration. The possible regulatory role of PAKs in cell metabolism has not been well explored, but increasing evidence suggests that a cell metabolic phenotype is related to cell interactions with the microenvironment. We analyzed the impact of PAK inhibition by small molecule inhibitors, small interfering RNA, or gene knockout on the rates of mitochondrial respiration and aerobic glycolysis. Pharmacological inhibition of PAK group I by IPA-3 induced a strong decrease in metabolic rates in human adherent cancer cell lines, leukemia/lymphoma cell lines, and primary leukemia cells. The immediate effect of FRAX597, which inhibits PAK kinase activity, was moderate, indicating that PAK nonkinase functions are essential for cell metabolism. Selective downregulation or deletion of PAK2 was associated with a shift toward oxidative phosphorylation. Alternatively, PAK1 knockout resulted in increased glycolysis. However, the overall metabolic capacity was not substantially reduced by PAK1 or PAK2 deletion, possibly due to partial redundancy in PAK1/PAK2 regulatory roles or to activation of other compensatory mechanisms.

## Introduction

P21-activated kinases (PAKs) participate in the regulation of virtually all processes associated with cytoskeletal rearrangements, such as cell migration, apoptosis, or cell division. The human PAK protein family consists of two groups, each of which contains the products of three different genes. Group I includes three serine/threonine kinases (PAK1 to PAK3), which have nearly identical kinase domains but at least partly different functions (Coniglio, Zavarella, & Symons, 2008; Grebenova et al., 2019; Kumar, Sanawar, Li, & Li, 2017). PAK3 expression is limited to a few cell types, whereas PAK1 and PAK2 are found in the majority of human cells.

PAK activity is often upregulated in cancer (Rane & Minden, 2019), and PAKs are therefore considered as potential therapy targets (Binder et al., 2019; Kanumuri et al., 2020; Senapedis, Crochiere, Baloglu, & Landesman, 2016). In relation to tumorigenesis, PAKs are involved in the modulation of cell adhesion and migration properties, which are critical for metastasis formation. Epithelial-to-mesenchymal transition (EMT), which promotes cancer cell invasiveness, was reportedly associated with changes in cell metabolism, another hallmark of oncogenic cell transformation. In general, tumor cells often rely on aerobic glycolysis (the so-called Warburg effect) to cover increased biosynthetic and energetic needs and to maintain the redox state. Nevertheless, they maintain the ability to switch among different metabolic phenotypes and to adapt to environmental changes (Kreuzaler, Panina, Segal, & Yuneva, 2020).

Increasing evidence points to functional associations between cell metabolism on the one hand and cell adhesion or migration on the other hand. Cell-cell and cell-matrix interactions have an impact on the cell metabolic phenotype, at least in solid tumors (Sousa, Pereira, & Paredes, 2019). Oxidative phosphorylation prevents cell migration by generating reactive oxygen species (ROS), which induces anoikis, a specific form of cell death caused by a loss of cell attachment (Lu, Tan, & Cai, 2015). In migrating cells, increased glycolysis is associated with deviation of intermediates to pentose phosphate pathway, which produces the major ROS scavenger nicotinamide dinucleotide phosphate (NADPH), providing protection from ROS-induced toxic effects (Sousa et al., 2019). In addition, metabolites produced by aerobic glycolysis promote EMT and cell migration (Han et al., 2013).

PAKs are considered key regulators of cell adhesion and migration, but several previous reports also indicated their involvement in the regulation of cell metabolism (Fig. 1). PAK2 was proposed to mediate E-cadherin signaling through activation of AMPK, a metabolic regulator promoting glucose uptake (Campbell, Salvi, O’Brien, Superfine, & DeMali, 2019). In muscle cells, insulin-induced glucose uptake is mediated by PAK1 (Tunduguru et al., 2014). PAK1 also enhances the activity of phosphoglucomutase 1, an enzyme that interconverts glucose-1-phosphate and glucose-6-phosphate (Gururaj, Barnes, Vadlamudi, & Kumar, 2004). Alternatively, PAK2 activity leads to decreased glucose uptake in neuronal cells (Varshney & Dey, 2016). Apart from glucose uptake and conversion, PAK2 is involved in a feedback loop activating the pyruvate kinase isoform PKM2, which supports pyruvate conversion to lactate (aerobic glycolysis) rather than the Krebs cycle supply (Cheng et al., 2018; Gupta et al., 2018). PAK1 activity is also connected to the kinase mTOR, which is a key metabolic regulator. However, the mTOR interactome is complex, and PAK1 was reported to be both upstream and downstream of mTOR, suggesting possible feedback loops (Kim et al., 2020; Rouquette-Jazdanian et al., 2015).

**Figure 1:**
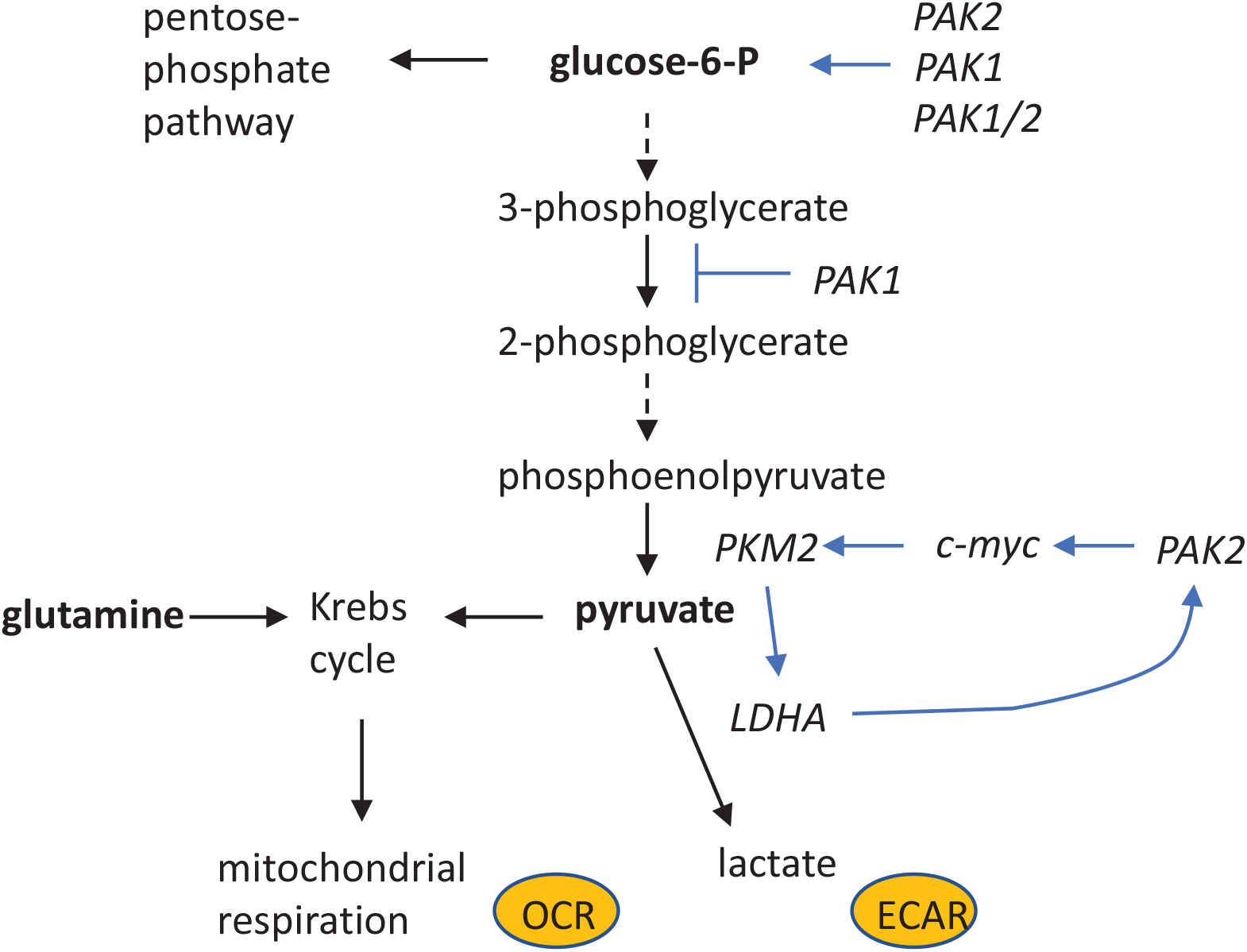
PAK involvement in the regulation of metabolic pathways. The scheme illustrates previously reported interaction points between PAK signaling and cell metabolism. In the present work, we analyzed the effect of PAK group I inhibition on the rates of oxidative phosphorylation (OCR) and lactate production (ECAR) in cancer/leukemia cell lines.

The current knowledge about different PAK functions is mainly based on adherent cell models. Cell adhesion processes are somewhat different in leukemia cells, which do not form stable cell-matrix or cell-cell contact points. Nevertheless, they interact with stromal cells, as well as with the extracellular matrix (ECM) of the bone marrow, and these interactions have a profound impact on their behavior. The adhesion structures of leukemia cells are more dynamic than those of adherent cells and they are usually not tightly connected to the actin cytoskeleton, but PAKs are still involved in the regulation of leukemia cell adhesion to the ECM (Kuželová et al., 2021). In addition, PAKs are indispensable for leukemia cell survival, as their inhibition leads to rapid cell death (Kuzelova, Grebenova, Holoubek, Roselova, & Obr, 2014; Kuželová et al., 2021; Pandolfi et al., 2015). Similar to solid tumors, leukemia cells often display a preference for aerobic glycolysis (Jiang & Nakada, 2016), and the metabolic phenotype has been proposed as a prognostic parameter for clinical outcome in acute myeloid leukemia (Chen, W. L. et al., 2014; Herst, Howman, Neeson, Berridge, & Ritchie, 2011). In leukocytes, PAK1 was reported to inhibit glycolysis by binding to phosphoglycerate mutase B, which converts 3-phosphoglycerate to 2-phosphoglycerate (Shalom-Barak & Knaus, 2002).

The emerging interconnection between adhesion and metabolic signaling, together with the key role of PAK in the regulation of cell adhesion and migration, prompted us to analyze the possible regulatory function of PAK group I in metabolic processes. In the present work, we modulated PAK activity in adherent and leukemia cells using several approaches: pharmacological inhibition of kinase or nonkinase PAK function, reduction of PAK transcript levels using siRNA, and gene knockout. The induced changes in the rates of mitochondrial respiration (OCR) and lactate production (ECAR) were monitored using the Seahorse XFp analyzer. The effect of inhibitors on glucose uptake was analyzed using a fluorescent derivative of 2-deoxyglucose, which was detected by flow cytometry.

## Materials & methods

### Cell lines and primary cells

HeLa, HEK293T, and HEL cells were obtained as a gift and authenticated using analysis of short tandem repeats. The results were compared with the ATCC database. OCI-AML3, Karpas-299, and Jurkat cell lines were purchased from DSMZ (Germany). After receipt, the cells were briefly expanded, and aliquots were cryopreserved for later use. The cell line MOLM-7 (not commercially available) was obtained from the laboratory of origin (Tsuji-Takayama et al., 1994). The majority of cell lines were cultured in RPMI-1640 medium with 10% fetal calf serum (FCS), 100 U/ml penicillin, and 100 µg/ml streptomycin at 37 °C in a 5% CO_2_ humidified atmosphere, except for HEK293T (DMEM medium) and OCI-AML3 (alpha-MEM medium, 20% FCS) cells.

Primary cells from patients with acute myeloid leukemia were obtained from leukapheresis at diagnosis. All patients provided written informed consent for the use of their biological material for research purposes, in accordance with the Helsinki Declaration. The project was approved by the Ethics Committee of the Institute of Hematology, and all experiments were performed in accordance with relevant guidelines and regulations. The leukapheretic products were diluted 20-fold in phosphate buffered saline (PBS), and the mononuclear cell fraction was then separated using Histopaque-1077 (Sigma, #H8889).

### Cell transfection

ON-TARGETplus siRNA (Dharmacon) targeting PAK1 (#L-003521) or PAK2 (#L-003597), in parallel with a nontargeting control (#D-001810), was transfected using jetPRIME transfection reagent (Polyplus Transfection) following the manufacturer’s instructions. The concentration range for siRNA was 100 to 150 nM (final concentration during transfection). The same protocol was used to transfect HeLa cell clones with the PAK2-GFP plasmid (Grebenova et al., 2019). The cells were then cultivated for 24 h, resp. 4 h (siRNA, resp. plasmid) without changing the medium, harvested, and seeded into Seahorse plates for cell metabolism measurement or into Petri dishes for protein analysis by western blotting.

### Gene knockout

To generate cells with PAK1 or PAK2 knockout via nonhomologous end-joining, we designed single guide RNA (sgRNA) sequences as follows ((N)_19-20_NGG strand, 5’ to 3’)): PAK1 sgRNA – ATTTGATGTCTGAAGCAAGC, PAK2 sgRNA – CATGTCTGATAACGGAGAAC, HPRT1 sgRNA -AAGTAATTCACTTACAGTC. Cas9 enzyme and sgRNA were obtained from IDT (Alt-R^®^ S.p. HiFi Cas9 Nuclease V3, Alt-R^®^ CRISPR–Cas9 sgRNA). HeLa cells were transfected with the enzyme and with PAK1 or PAK2 sgRNA using nucleofection (4D-Nucleofector™ X Unit, Lonza). Due to low transfection efficiency in leukemia cell lines, OCI-AML3 cells were cotransfected with sgRNA targeting the hypoxanthine phosphoribosyltransferase 1 (HPRT1) gene, allowing for enrichment of successfully transfected cells through resistance to 6-thioguanine (6-TG). The typical transfection reaction contained 4×10^5^ HeLa cells or 1×10^6^ OCI-AML3 cells, 500 pmol Cas9 enzyme, and 600 pmol sgRNA (2×300 pmol in the case of HPRT1 cotransfection). After transfection, HeLa cells were seeded in 6-well plates and grown according to the experimental settings. OCI-AML3 cells were seeded in 6-well plates, 6-TG was added (10 µg/ml final concentration) after 3 days, and the cells were cultured for at least 10 days with regular replacement in medium containing 6-TG.

### Limiting dilution cloning

To obtain clones stably modified with PAK2 knockout, a Hela cell suspension was diluted to 5 cells/ml 3 days after CRISPR/Cas9 modification, and 100 µl aliquots were distributed into a 96-well culture plate. Cell growth was regularly monitored by visual inspection, and wells containing a single colony were selected for further cell expansion and analysis.

### Inhibitors and antibodies

IPA-3 (#3622) and PIR3.5 (#4212) were purchased from Tocris Bioscience and dissolved in sterile dimethylsulfoxide (DMSO) to make 50 mM stock solutions. Working solutions were prepared by 10-fold dilution of the stock solution in 50 mM Tris, pH 8.0, immediately before use. FRAX597 (#6029) was purchased from Tocris Biosciences and dissolved in sterile DMSO to make a 10 mM stock solution. Working solution was prepared by 10-fold dilution in cell culture medium. Dasatinib was obtained from Selleckchem (S1021), and a 200 µM stock solution was made in sterile DMSO. The following antibodies were used: PAK1 (Abcam, ab223849), PAK2 (Abcam, ab76293), PAK1 (phosphoS144)+PAK2 (phosphoS141)+PAK3 (phosphoS139) (Abcam, ab40795), PAK1/2/3 (Cell Signaling, #2604), and β-actin (Sigma–Aldrich, A5441).

### Cell metabolism measurement

The oxygen consumption rate (OCR) and the extracellular acidification rate (ECAR) were measured using the MitoStress Test kit (Agilent, #103010–100) following the manufacturer’s instructions. The last injection was supplemented with 2-deoxyglucose (2-DG, Sigma, D8375) to obtain the ECAR background value due to nonglycolytic acidification. If not otherwise indicated, the final well concentrations were as follows: oligomycin (OM) 1 µM, carbonyl cyanide-4 (trifluoromethoxy) phenylhydrazone (FCCP) 0.3 and 0.5 µM, rotenone/antimycin A 0.5 µM, and 2-DG 50 mM. In the experiments involving glycolytic stress, glucose (10 mM) and pyruvate (1 mM) were omitted from the Seahorse medium, and the injections were as follows: glucose 10 mM, OM 1 µM, FCCP 0.4 µM, and rotenone/antimycin A 0.5 µM with 50 mM 2-DG. CellTak (Corning, #354240) was used to coat Seahorse XFp plates according to the manufacturer’s instructions for all leukemia cells, as well as for HEK293T cells. HeLa (16,000 cells/well) or HEK293T (20,000 cells/well) cells were seeded in 80 µl culture medium and incubated overnight. The plate was then washed with Seahorse medium (Agilent #103676–100 or #103575–100, respectively), and the cells were treated with inhibitors if appropriate and incubated for 1 h at 37 °C without CO_2_. The control cells were treated with the solvent only. When IPA-3 and PIR3.5 were used for cell treatment, the plates were washed twice with Seahorse medium and analyzed using the Seahorse XFp apparatus (Agilent). Cells treated with FRAX597 or transfected cells were measured without washing. After measurement, the medium was aspirated, the plate was cryopreserved, and the relative cell numbers in the individual wells were assessed using the CyQuant Cell Proliferation Assay Kit (Molecular Probes, #C7026). Seahorse records were corrected for well-to-well differences in the cell numbers and analyzed using Wave software (Agilent). Leukemia cells (40,000 cells/well for cell lines, 150,000–200,000 cells/well for primary cells) were seeded in 50 µl Seahorse medium (Agilent #103676–100), briefly centrifuged (1 min, 200 g, gentle brake), incubated for 25 min at 37 °C without CO_2_ prior to the addition of 130 µl medium, treated with inhibitors for 1 h, and subjected to analysis.

### Glucose uptake assay

Cells (5×10^5^) were washed once in PBS, resuspended in 0.5 ml Seahorse medium with glutamine without glucose, and treated with the inhibitors for 1 h. A fluorescent derivative of 2-deoxyglucose (2-NBDG, Invitrogen, N13195) was added at a final concentration of 20 µM for a further 30 min incubation at 37 °C. Then, the cells were washed in cold PBS (300 g/4 °C/5 min) and resuspended in 0.5 ml cold PBS. The samples were placed on ice, and 2-NBDG fluorescence was immediately analyzed using the FITC channel on a BD Fortessa flow cytometer. Propidium iodide (10 µg/ml final concentration) was added before measurement to exclude dead cells from the analysis, and the cell debris was outgated in scattergrams. The mean fluorescence intensity (MFI) of control cell samples without 2-NBDG was used to determine the background, which was subtracted from the corresponding MFI of samples with 2-NBDG.

### Western blot

Adherent cells were washed once with ice-cold HBS (HEPES – buffered saline; 20 mM HEPES, 150 mM NaCl, pH 7.1) and scrapped into Pierce IP Lysis Buffer (#87787) with freshly added protease and phosphatase inhibitors. The suspension was then transferred to a centrifugation tube and incubated for 10 min at 4 °C. Cellular debris was removed by centrifugation (15,000 g/4 °C/15 min), and the lysate was mixed 1:1 (v/v) with 2× Laemmli sample buffer and incubated for 5 min at 95 °C. An equivalent of 20 µg total protein was resolved on a 7.5% polyacrylamide gel (10×6 cm or 18×18 cm) and transferred to a nitrocellulose membrane. The membrane was blocked for 1 h in 3% bovine serum albumin and incubated for 1 h with the primary antibody in PBS with 0.1% Tween-20 (PBST) at room temperature. Thereafter, the membrane was washed in PBST six times and incubated with the corresponding HRP-conjugated secondary antibody for 1 h. The chemiluminescence signal from Clarity Western ECL Substrate (Bio-Rad, #170–5060) was detected and analyzed using G:BOX iChemi XT-4 (Syngene).

### Statistical analyses

Statistical evaluation of the experimental results was performed using GraphPad Prism version 7.03 (GraphPad software, San Diego, California USA). The p value limit for statistically significant differences between groups was set to 0.05.

## Results

### Study design

Mitochondrial respiration and aerobic glycolysis (lactate production) were analyzed using the Seahorse XFp device, which monitors the oxygen consumption rate (OCR) in parallel with the extracellular acidification rate (ECAR). An example of a Seahorse XFp record is shown in Supplementary Fig. S1, which also depicts the basal and maximal metabolic rates. The standard Seahorse assay was performed in serum-free culture medium supplemented with glucose, pyruvate, and glutamine. After signal stabilization (4 cycles), three compounds were added in four injections: oligomycin (OM) inhibits ATP synthase, protonophore FCCP (two injections) dissipates the proton gradient across the mitochondrial membrane, and a mixture of rotenone with antimycin A and 2-deoxyglucose (rot/A+2-DG) inhibits both the respiration chain and glycolysis, thereby giving the signal background deduced from both the basal and maximal metabolic rates. To determine the effect of PAK inhibition on processes not associated with glucose uptake and/or consumption, OCR was also measured in glucose-free and pyruvate-free medium. Under these conditions, the ECAR values were very low, and the cells mainly relied on the oxidative phosphorylation to produce energy.

The amount and/or activity of PAK were modified using small molecule inhibitors, small interfering RNAs (siRNAs), or CRISPR/Cas9 knockout. The inhibitor IPA-3 binds to inactive PAK group I molecules and prevents conformational changes that are necessary for PAK activation (Deacon et al., 2008). A structural isomer of IPA-3, called PIR3.5, displays no inhibitory activity toward PAK1 (Deacon et al., 2008) and is thus commonly used as a negative control compound to IPA-3. An alternative PAK inhibitor, FRAX597, binds to the ATP-binding site of kinase-active PAK molecules and prevents PAK kinase activity but presumably does not directly interfere with nonkinase PAK functions (Licciulli et al., 2013). We have previously described the effects of these inhibitors on cell adhesion and viability in adherent HeLa and HEK293T human cancer cell lines (Grebenova et al., 2019) as well as in human leukemia cells (Kuželová et al., 2021). In the present work, we used PAK inhibitors at previously optimized concentrations regarding efficiency and toxicity: IPA-3 and PIR3.5 were added at 20 and 50 µM, and FRAX597 was added at 2 and 10 µM. The efficiency of PAK silencing by siRNA or by gene knockout was assessed by western blotting using previously characterized antibodies against PAK1, PAK2, or the autophosphorylated site Ser144/141 of PAK1/PAK2 (Grebenova et al., 2019).

### Effect of PAK inhibitors on cell metabolism in adherent cells

HeLa or HEK293T cells were seeded into Seahorse plates and cultured for 24 h. The medium was replaced by the corresponding serum-free variant, and the cells were treated for 1 h with inhibitors. Treatment with IPA-3 substantially reduced both the OCR and ECAR (Fig. 2A). The effect of the control compound PIR3.5 was more moderate, if any. In contrast to IPA-3, only limited changes in the cell metabolic rates were observed after 1 h FRAX597 treatment, which only inhibited the kinase function of PAK (Fig. 2B).

**Figure 2:**
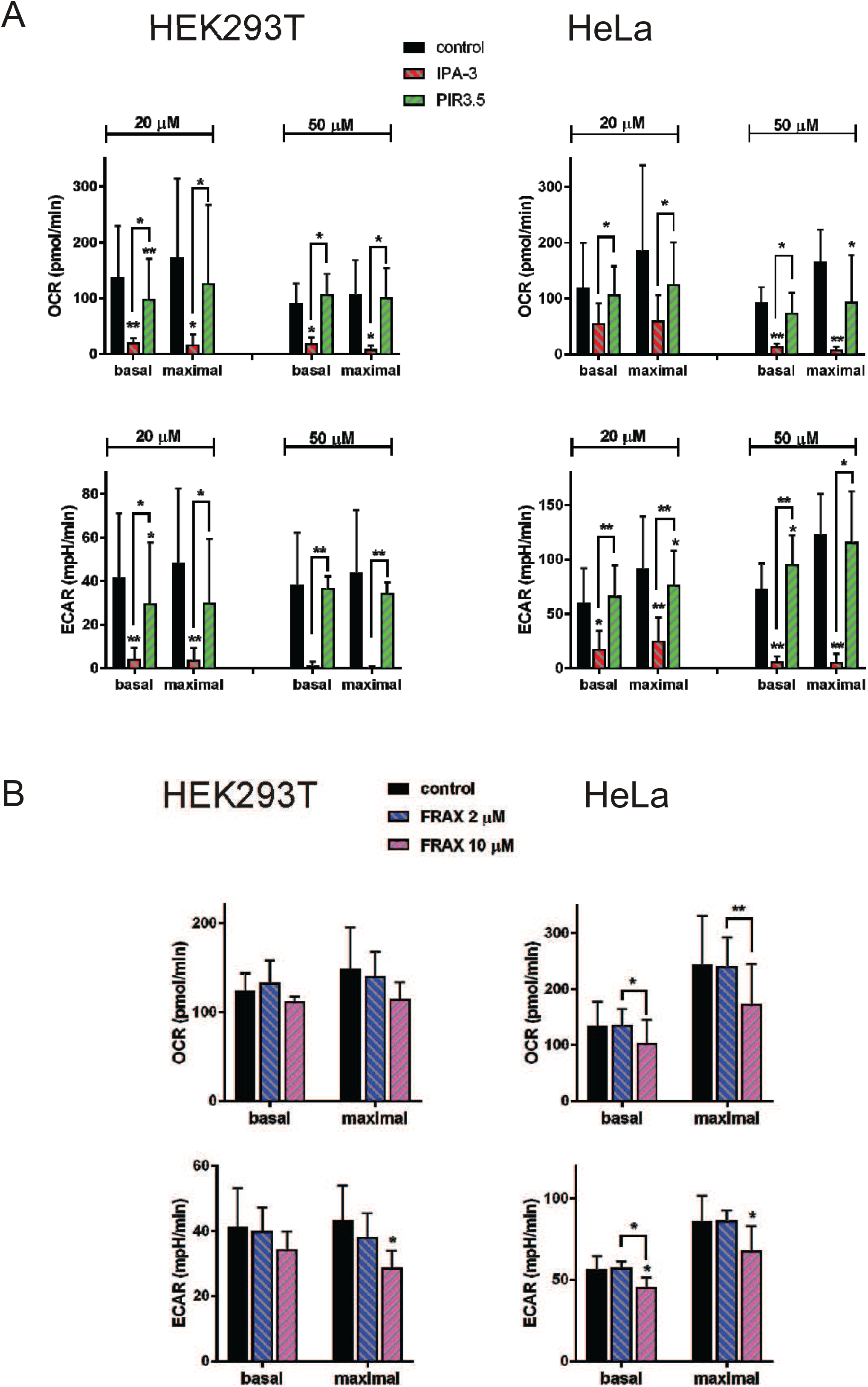
Effect of inhibitors on metabolic rates in adherent cells. Adherent HEK293T (left) or HeLa (right) cells were treated with inhibitors in duplicate and incubated for 1 h at 37 °C without CO_2_ prior to metabolic measurement. The oxygen consumption rate (OCR) and the extracellular acidification rate (ECAR) were determined from Seahorse XFp records, as shown in Supplementary Fig. S1. The records were first corrected to well-to-well differences in the cell numbers, and duplicates were averaged. The graphs show the means and standard deviations of basal and maximal metabolic rates obtained from repeated experiments. Differences between treated samples and untreated controls were evaluated using standard paired Student’s t-test. * p < 0.05, ** p < 0.01. A. Effect of IPA-3 (red bars) and PIR3.5 (green bars). Three to seven independent experiments were performed for each cell line and for each inhibitor concentration (20 or 50 µM as indicated). B. Effect of FRAX597 at 2 µM (blue bars) or 10 µM (magenta) concentration. Four independent experiments were performed for each cell line.

### Effect of PAK inhibitors on cell metabolism in leukemia cells

Leukemia cells are reportedly very sensitive to PAK inhibition (Kuzelova et al., 2014; Kuželová et al., 2021; Pandolfi et al., 2015), and treatment with IPA-3 induces cell death in the majority of leukemia cell lines. Accordingly, both the OCR and ECAR were markedly reduced by 20 µM IPA-3 in all leukemia cell lines tested (Fig. 3A), except for MOLM-7, which has a very low amount of PAK1 and a high PAK2 level (Kuželová et al., 2021). Increasing the IPA-3 dose to 50 µM was sufficient to achieve a marked decrease in the OCR and ECAR also in the latter cell line (data not shown). Consistent with the findings in adherent cell models (Fig. 2), inhibition of PAK kinase activity by FRAX597 had a lower impact on the metabolic rates than treatment with IPA-3 within the first 1 h (Fig. 3B versus 3A). Nevertheless, leukemia cell metabolic preference was slightly shifted to glycolysis by FRAX597 treatment. In addition, longer treatment (5 h) with FRAX597 resulted in a large decrease in the OCR and ECAR values similar to that induced by IPA-3 (Supplementary Fig. S2). We also performed several experiments with primary cells from patients with acute myeloid leukemia (Supplementary Fig. S3). As the signal from these cells was lower than that from cell lines, a higher number of cells (150 to 200 thousand per well) was required for Seahorse XFp measurement compared to the cell lines (40 thousand per well). We found previously that the efficiency of IPA-3 strongly depends on the cell density. Therefore, a primary cell suspension at 3×10^5^ cells/ml was pretreated with IPA-3 or PIR3.5 for 1 h, and the cells were washed and seeded into Seahorse plates for metabolic measurement. Under these conditions, the effects of 20 µM IPA-3 were not consistent (Fig. S3A). However, the higher dose (50 µM) induced similar changes as in other cells, i.e., marked reduction of both OCR and ECAR (Fig. S3B).

**Figure 3:**
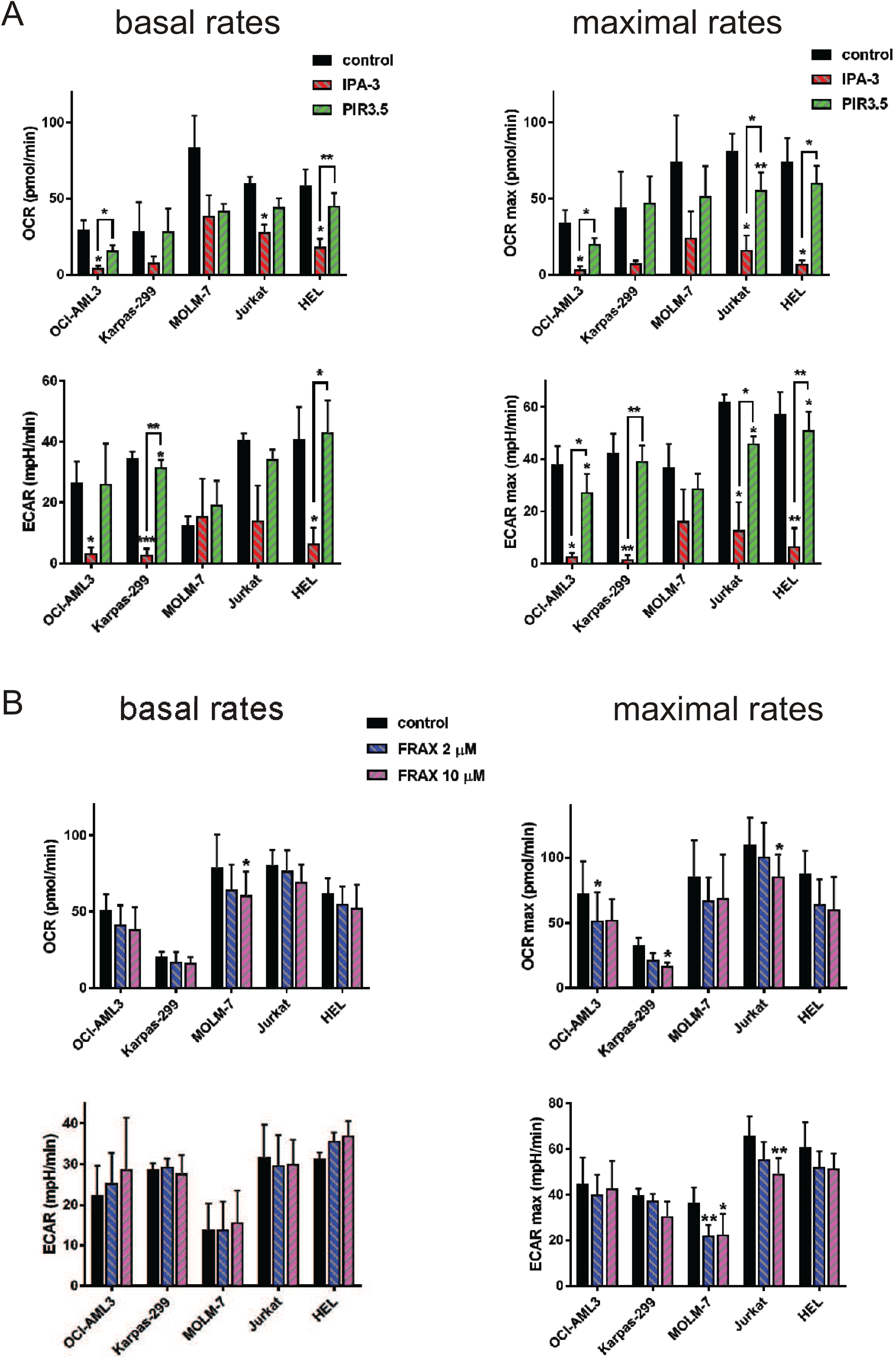
Effect of inhibitors on metabolic rates in leukemia cell lines. Cells (40,000 per well) were seeded into CellTak-coated wells and incubated for 25 min prior to 1 h treatment with inhibitors. The graphs show the means and standard deviations of basal (left) and maximal (right) metabolic rates from repeated experiments. Three independent experiments were performed for each condition, and differences between treated samples and untreated controls were evaluated using standard paired Student’s t-test. * p < 0.05, ** p < 0.01. A. Effect of 20 µM IPA-3 (red bars) or PIR3.5 (green bars). B. Effect of FRAX597 at 2 µM (blue bars) or 10 µM (magenta) concentration.

### Effect of PAK inhibition on glucose uptake

Previous reports indicated PAK involvement in glucose uptake in some cell types, and limited glucose supply could explain the observed reduction of both OCR and ECAR in our model systems. We thus searched for possible involvement of PAK in the glucose uptake rate using a fluorescent derivative of 2-deoxyglucose (2-NBDG). The impact of IPA-3 pretreatment was cell type-specific, but no systematic reduction in glucose uptake was found (Fig. 4A). Interestingly, glucose uptake was enhanced by the control compound PIR3.5 in some cell lines.

**Figure 4:**
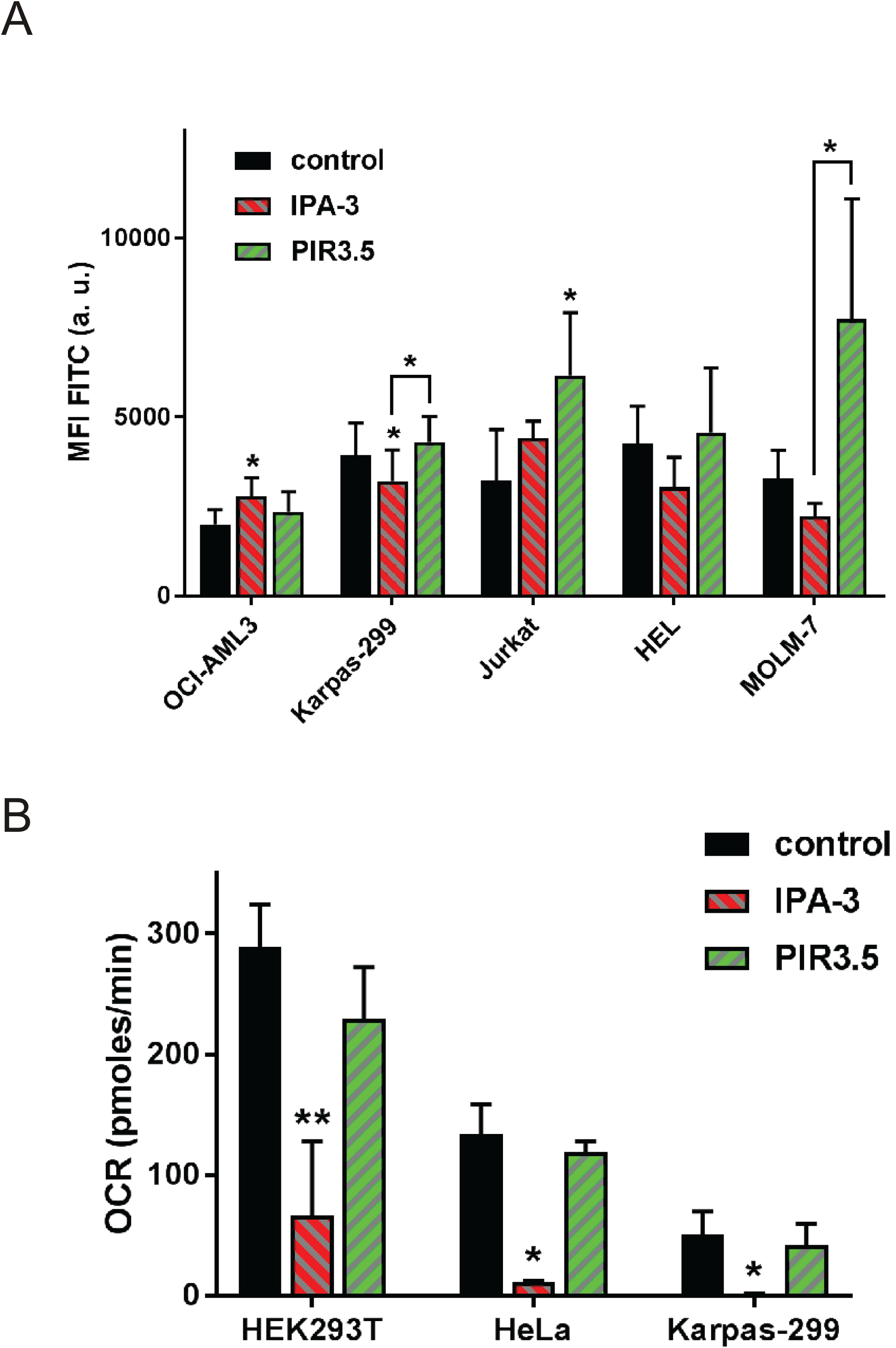
Effect of IPA-3 on glucose uptake and on cell respiration under glycolytic stress. A. The cells were seeded in glucose/pyruvate-free medium and treated with 20 µM IPA-3 or PIR3.5 for 1 h prior to the addition of 20 µM 2-NBDG. After a further 30 min incubation, the cells were washed and the amount of accumulated 2-NBDG was determined by flow cytometry. The background mean fluorescence intensity (MFI) was obtained from control tubes without 2-NBDG and subtracted. The bars represent the means and standard deviations from four independent experiments for each cell line. The differences between the treated samples and the corresponding controls were evaluated using standard paired Student’s t-test. * p < 0.05. B. Basal metabolic rates were measured in the absence of glucose and pyruvate. ECAR values were very low under these conditions (see examples of Seahorse records in the Supplementary Fig. S4). The cells were seeded in glucose/pyruvate-free medium and incubated for 1 h in the presence of 20 µM IPA-3 or PIR3.5 prior to measurement. The graph represents the means and standard deviations from three independent experiments for each cell line. The differences between the treated samples and the corresponding controls were evaluated using standard paired Student’s t-test. * p < 0.05, ** p < 0.01

### Effect of PAK inhibition under glycolytic stress conditions

PAK2 reportedly supports aerobic glycolysis rather than the Krebs cycle supply and oxidative phosphorylation (Fig. 1). However, the observed large reduction in both OCR and ECAR upon PAK inhibition by IPA-3 indicated that PAK might also be required for mitochondrial respiration. We thus performed Seahorse measurements in the absence of glucose and pyruvate, thus limiting the use of the glycolytic pathway. Examples of records are provided in Supplementary Figure S4. The measured OCR values from these experiments are summarized in Fig. 4B, showing that the strong effect of IPA-3 was retained under these conditions.

### Effect of siRNA-mediated PAK silencing

Small interfering RNAs (siRNAs) were used to discriminate between PAK1 and PAK2 functions in cell metabolism. The efficiency of silencing was measured by western blotting, and examples are given in Supplementary Fig. S5. As summarized in Fig. 5A, we observed some cross-effects at the PAK protein level: siRNA PAK1 reduced PAK2 protein levels in HEK293T cells (left), and conversely, siRNA PAK2 reduced PAK1 protein levels in both HeLa and HEK293T cells (right), which might be due to mutual interaction and possibly to mutual regulation between PAK1 and PAK2 (Grebenova et al., 2019). HeLa cells have low PAK1 levels compared to HEK293T cells (Grebenova et al., 2019). Accordingly, PAK1 targeting by siRNA PAK1 was more efficient in HeLa cells (black bars in Fig. 5A, left).

**Figure 5:**
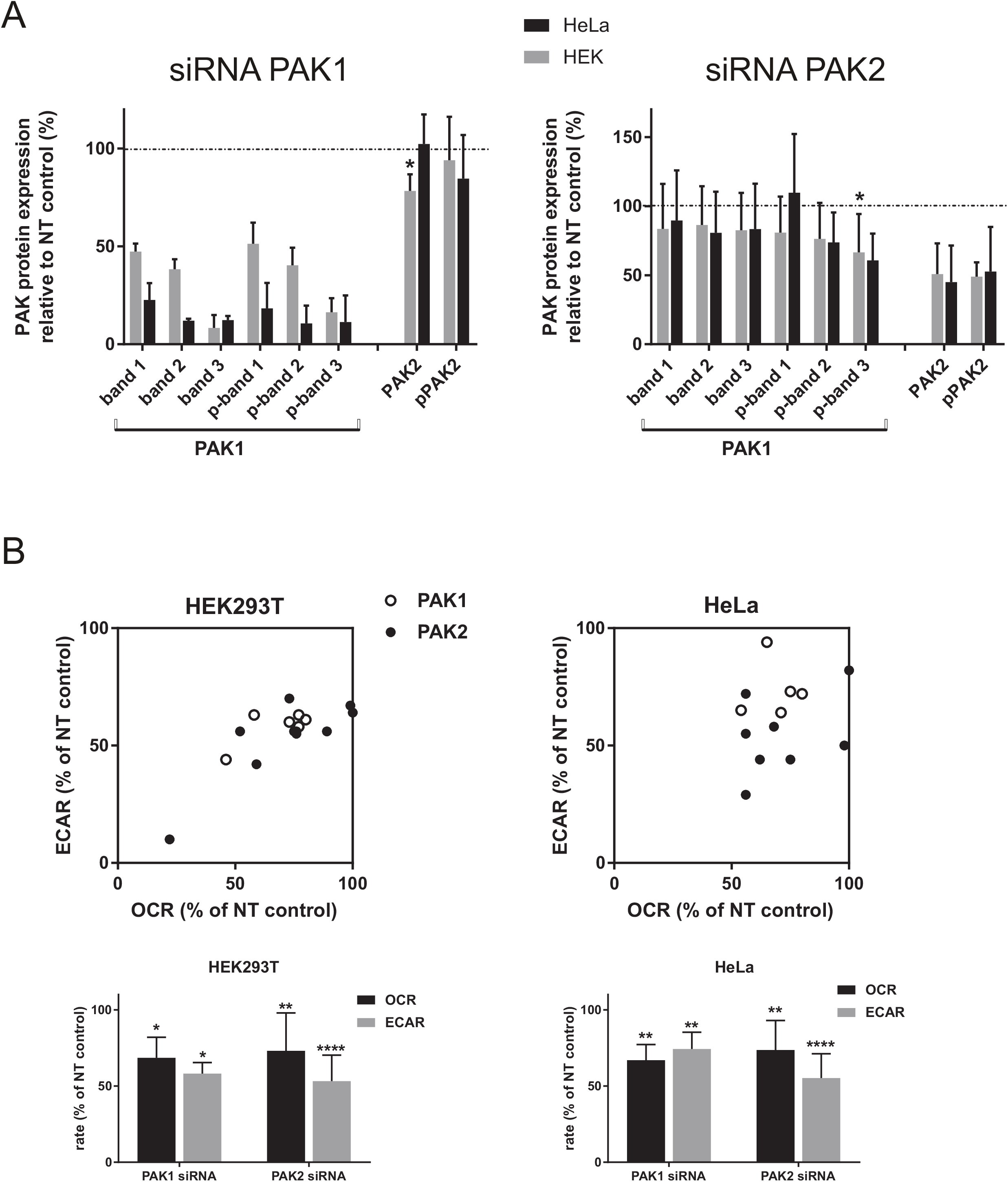
Effect of siRNA targeting PAK1/PAK2. The cells were transfected with nontargeting (NT), PAK1, or PAK2 siRNA and cultured for 24 h under standard conditions. Then, they were harvested, and aliquots were seeded into microtitration plates in triplicate (NT siRNA and PAK1/PAK2 siRNA) and cultured for additional 24 h prior to analysis on Seahorse XFp. The remaining cells were further cultured for 24 h in Petri dishes and used to assess PAK1/PAK2 expression levels and PAK phosphorylation at Ser144/141 by western blotting. A. Reduction of PAK levels by siRNA. Western blot band intensity values were expressed relative to the corresponding NT siRNA controls. The labeling of individual PAK1 bands is shown in Supplementary Figure S5. The graphs show mean values and standard deviations from three independent experiments for the HeLa cell line and from three, resp. six experiments for HEK293T cells transfected with siRNA PAK1, resp. PAK2. In HEK293T cells, the cross-effects of PAK1 siRNA on PAK2 levels and of PAK2 siRNA on PAK1 p-band 3 were statistically significant (* p < 0.05). B. The measured OCR and ECAR values were corrected for differences in cell numbers, the triplicates were averaged, and the rates were expressed as relative to the corresponding NT siRNA control. Top: Correlation of changes in OCR and ECAR in individual experiments. Bottom: The graphs show the means and standard deviations of metabolic rates obtained from repeated experiments. The differences between samples transfected with PAK siRNA and the corresponding NT controls were evaluated using standard paired Student’s t-test. * p < 0.05, ** p < 0.01, *** p < 0.001, **** p < 0.0001.

Fig. 5B shows the results of individual Seahorse XFp experiments involving cells treated with PAK1 or PAK2 siRNA (top) as well as the summary and statistical evaluation of these results (bottom). Both PAK1 and PAK2 siRNA reduced both the OCR and ECAR, and the effect of PAK2 siRNA was more pronounced on the ECAR values. Treatment with siRNA also resulted in a decrease in the OCR in the absence of glucose and pyruvate (Supplementary Fig. S6).

### Effect of PAK knockout

As an alternative method to specifically reduce PAK1 or PAK2 expression, we performed *PAK1/PAK2* gene knockout (KO) in HeLa cells using the CRISPR/Cas9 system. Western blot analysis confirmed a substantial reduction in the respective PAK1/PAK2 protein levels (Fig. 6A). Three different antibodies were used to detect PAK2. One of them, denoted as PAK2 (N-term), targets an epitope within the first 100 amino acids of PAK2 according to the manufacturer’s datasheet. The second antibody (PAK2) recognizes the carboxy-terminus, and its affinity for PAK is thus not modified by DNA repair occurring after CRISPR-induced DNA break. The third antibody recognizes the autophosphorylation site at Ser141, and the signal (pPAK2) thus corresponds to kinase-active PAK2. The summary graph in Fig. 6A shows that both PAK1 and PAK2 protein levels were consistently reduced to approximately 20 % in repeated experiments with CRISPR/Cas9 modifications. Interestingly, the decrease in PAK2 was virtually higher using the antibody targeting the N-terminus. This suggests that the antibody binds to the region affected by the DNA break induced by CRISPR/Cas9 and that the efficiency of PAK2 targeting is very high. While subsequent repair presumably leads to a frameshift-inducing mutation in the majority of cells, some cells can produce a modified but still functional protein. The impact of PAK KO on HeLa cell metabolic rates is shown in Fig. 6B. The overall metabolic activity of cells with PAK1 or PAK2 KO was not much different from that of the control cells (nucleofection without CRISPR/Cas9 modification). However, the metabolic phenotype was shifted toward glycolysis in PAK1 KO and toward oxidative phosphorylation in PAK2 KO.

**Figure 6:**
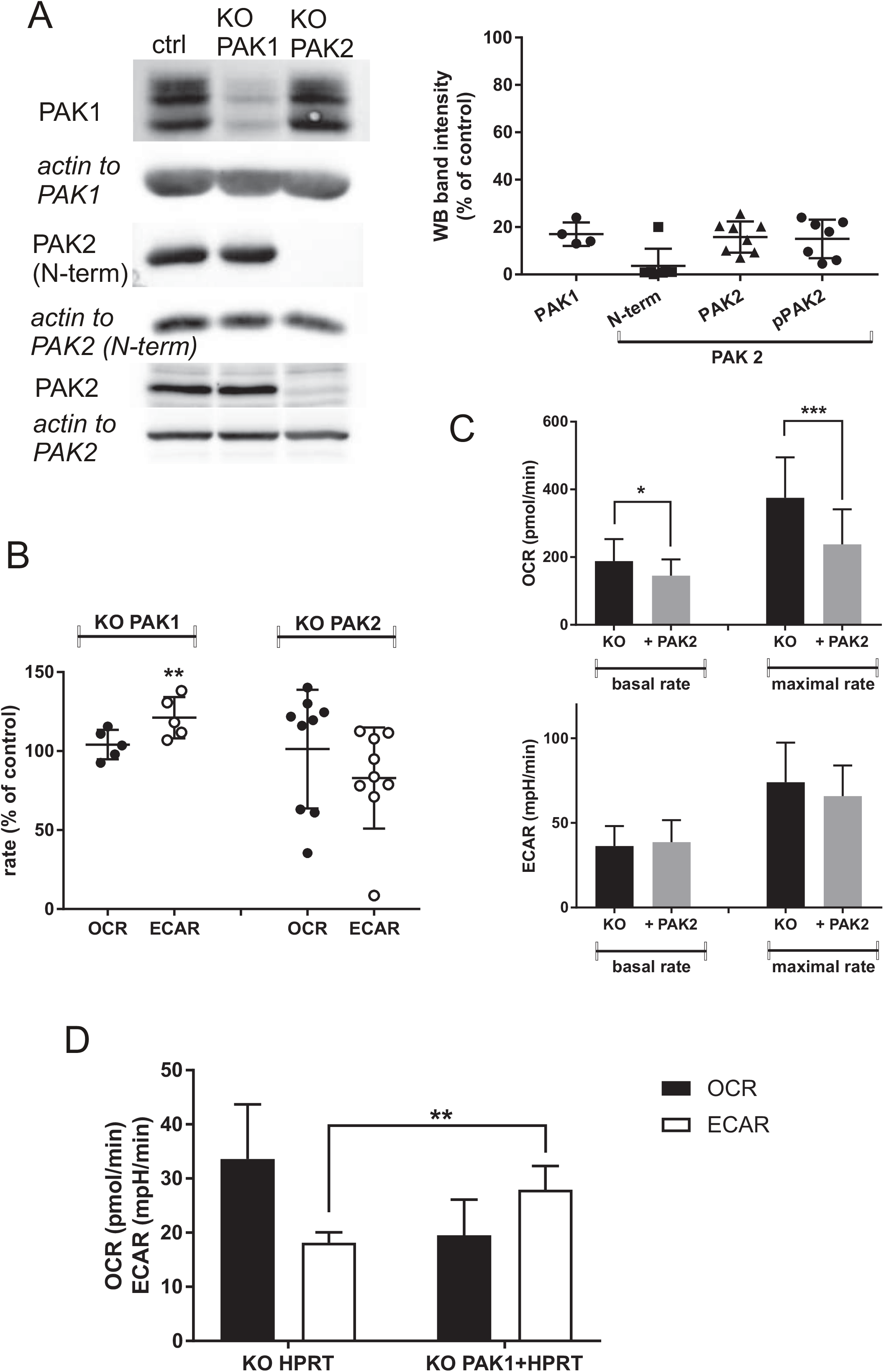
Effects of CRISPR/Cas9 PAK knockout. A. HeLa cells were subjected to CRISPR/Cas9 knockout (KO) of PAK1 or PAK2, and PAK protein levels were assessed by western blot 3 to 11 days later. The panel shows representative western blots and a summary of relative band intensities from repeated experiments. B. Metabolic rates of HeLa cells with PAK1 or PAK2 KO were measured using Seahorse XFp and expressed as relative to the values from control cells (empty nucleofection). The statistical analysis of differences between the cells with PAK KO and the corresponding controls was performed using the absolute values of OCR/ECAR (** p < 0.01). C. Clone D3 obtained from HeLa cells after PAK2 KO was transfected with a plasmid for PAK2-GFP expression, and the metabolic rates were measured using Seahorse XFp. Summary results from 4 independent transfections. The difference between PAK2-GFP-expressing cells (denoted as + PAK2) and the corresponding control (KO) was evaluated using the standard paired Student’s t-test (* p < 0.05, *** p < 0.001). D. In OCI-AML3 cells, the HPRT gene was targeted simultaneously with PAK1 by the CRISPR/Cas9 system, and the modified cell population was subsequently enriched through 6-thioguanine selection. Metabolic rates were compared between cells with double PAK1+HPRT KO and cells with single HPRT KO. The bars show the means and standard deviations from four experiments.

The results from repeated experiments with PAK2 KO were less conclusive than those with PAK1 KO, and we thus attempted to establish stably modified HeLa clones by the limiting dilution method. Of the 12 expanded clones, 11 had a low-to-undetectable signal from the PAK2 (N-term) antibody (examples are given in Supplementary Fig. S7A) but still produced PAK2, as detected by the antibody recognizing the C-terminus, as well as by the phospho-specific antibody targeting pSer141. We also noted variability in the western blot band position, indicating slightly different lengths of pPAK2 products in the individual clones (Fig. S7A). Two clones with reduced PAK2/pPAK2 levels were selected to analyze the effect of PAK2 re-expression. These clones were transfected with a previously described plasmid designed for exogenous expression of PAK2 tagged with green fluorescent protein (PAK2-GFP) (Grebenova et al., 2019). The efficiency of transfection was assessed by flow cytometry and found to be 40% and 31% on average in clones D3 and D12, respectively. Examples of flow cytometry dotplots are given in Supplementary Fig. S7B. Fig. 6C shows the summary results of metabolic analyses performed with clone D3. Despite the limited transfection efficiency, PAK2 overexpression was associated with a decrease in the respiration rate (both basal and maximal OCR). The results obtained with clone D12 showed similar trends (Supplementary Fig. S7C) but without statistical significance, possibly due to the lower transfection efficiency in this clone.

The CRISPR/Cas9-based approach was also applied to leukemia cell lines, but the efficiency of gene modification was much lower in these cells. We thus used simultaneous HPRT KO, which provides resistance to 6-thioguanine, to enrich the OCI-AML3 population with PAK KO. The cells were nucleofected with guide RNA (gRNA) for HPRT (control sample) or with a mixture of gRNA for HPRT and PAK1 (test sample), and the cells were cultured in the presence of 6-thioguanine. Compared to cells with single HPRT KO, OCI-AML3 with PAK1 KO had reduced OCR and increased ECAR (Fig. 6D). Unfortunately, attempts to reduce PAK2 expression by gene KO in OCI-AML3 cells were not successful, possibly because PAK2 is essential for leukemia cell survival.

### Effect of dasatinib and PIR3.5/FRAX597 combination

PAK1 is an important downstream effector of c-Src in adherent cells. Therefore, we also looked for a possible effect of Src kinase inhibition by dasatinib on the metabolic rates in HeLa and HEK293T cells. Supplementary Fig. S8 shows that dasatinib induced no significant metabolic change.

The milder effect of FRAX597 compared to IPA-3 indicated that nonkinase activity of PAK is essential for cell metabolism. However, as the control compound PIR3.5 also affected cell metabolism, IPA-3 might have unspecific effects, such as reactive oxygen species generation, which could act in synergy with inhibition of PAK kinase activity. To check for this possibility, we analyzed the effect of combined treatment with FRAX597 and PIR3.5. However, we detected no synergy between these two compounds (Supplementary Fig. S9).

## Discussion

Our results suggest that PAK group I regulates the cell metabolism in a variety of cell types. Treatment with IPA-3, an allosteric inhibitor considered specific for PAK group I, rapidly inhibited both cell respiration and aerobic glycolysis in adherent cancer cell lines (Fig. 2A), leukemia cell lines (Fig. 3A), and primary leukemia cells (Supplementary Fig. S3). In primary cells, higher IPA-3 doses (50 µM instead of 20 µM) were needed to obtain consistent reduction of OCR and ECAR, but this might be due to different protocols used for treatment because of the higher cell density required for Seahorse XFp measurement. Nevertheless, we cannot exclude a lower sensitivity of primary cells to PAK inhibition compared with rapidly proliferating established cell lines. Additionally, MOLM-7 cells were more resistant to IPA-3 treatment than other leukemia cell lines (Fig. 3A), which may be due to higher PAK2 content in the MOLM-7 cell line (Kuželová et al., 2021) because increasing the IPA-3 dose to 50 µM was sufficient to obtain similar effects as in other cell lines.

IPA-3 could have additional unspecific effects due to increased oxidative stress. Indeed, the control compound PIR3.5 also affected the OCR and ECAR (Figs 2A, 3A, and S3). However, these nonspecific effects were generally milder and varied according to cell type. For example, basal OCR reduction due to PIR3.5 treatment was compensated by increased glycolysis in HeLa cells (Fig. 2A). A similar effect was observed in one primary leukemia sample (Fig. S3B) but not in HEK293T cells, perhaps because of the low reserve glycolytic capacity: the maximal ECAR rate was close to the basal ECAR rate in the latter cell line.

Although FRAX597 is less specific and might inhibit other kinases in addition to PAK group I (Licciulli et al., 2013), it had only a limited effect on the metabolic rates within 1 h (Figs 2B and 3B). PAK kinase activity is rapidly and largely inhibited by FRAX597 at the doses used (Grebenova et al., 2019; Kuželová et al., 2021), and we thus suggest that a nonkinase PAK function is essential for cell metabolism. Possible synergy between targeting PAK kinase activity and nonspecific effects, which might have explained the observed strong effect of IPA-3, was ruled out by analysis of combined treatment with FRAX597 and PIR3.5 (Supplementary Fig. S9).

Pharmacological inhibition of PAK group I using IPA-3 is known to induce cell death, especially in leukemia cells (Kuželová 2014, Pandolfi 2015). We can only speculate if the observed cytotoxicity is directly due to the suppression of cell metabolism, but such a causal relation would be supported by the fact that the effect of IPA-3 on OCR/ECAR was rapid (within 1 h), whereas cell death symptoms start to appear in several hours (Kuželová et al., 2021). The abrupt disruption of both mitochondrial respiration and aerobic glycolysis presumably causes severe energetic deficiency, and cell death induction is an expected outcome under such conditions. Consistently, longer treatment (5 h) with FRAX597 also induced both a large decrease in the metabolic rates (Supplementary Fig. S2) and subsequent cell death (Kuželová et al., 2021).

Small interfering RNA targeting either PAK1 or PAK2 was used to discriminate between these PAK group I members. However, cross-effects were usually detected except for siRNA PAK1 in HeLa cells, where PAK2 levels were not altered (Fig. 5A). Therefore, CRISPR/Cas9 knockout of the PAK1 or PAK2 gene was performed in HeLa cells to further confirm the observed differences between PAK1 and PAK2 (Fig. 6). Regardless of the method used, reduced PAK2 levels were associated with a shift of metabolic preferences toward oxidative phosphorylation in both HeLa and HEK293T cells (Figs 5B and 6B). Additionally, PAK2 overexpression in HeLa clones with PAK2 KO had the reverse effect, i.e., a decrease in the basal OCR, but not in the basal ECAR, which was detectable despite limited transfection efficiency (only 40% of cells overexpressed PAK2 in the experiments presented in Fig. 6C). Therefore, PAK2 might be specifically involved in pyruvate conversion to lactate, consistent with previous reports indicating that PAK2 favors aerobic glycolysis to the detriment of the Krebs cycle and oxidative phosphorylation (Fig. 1).

Alternatively, PAK1 targeting in HeLa or OCI-AML3 cells resulted in a more glycolytic phenotype (Figs 5B, 6B, and 6C). These results support a previous report indicating that PAK1 inhibits glycolysis in leukocytes (Shalom-Barak & Knaus, 2002). In HEK293T cells, PAK1 silencing using siRNA was less efficient and induced a decrease in PAK2 levels (Fig. 5A), and the results are thus not conclusive. This cross-reduction in protein levels could be due to the interaction between PAK1 and PAK2 proteins (Grebenova et al., 2019) rather than to cross-targeting of transcripts by siRNA.

Whereas acute inhibition of both PAK1 and PAK2 by IPA-3 led to a strong reduction in the metabolic rates (Figs 2A, 3A, and S3), PAK transcript targeting by siRNA induced more moderate changes (Fig. 5B). It is possible that the residual PAK amount in cells treated with siRNA was sufficient to maintain cell metabolism. Alternatively, PAK1 and PAK2 regulatory roles might be partly redundant. Indeed, the PAK1 protein translated from the dominant PAK1 transcript isoform (PAK1Δ15) is similar to PAK2 (Grebenova et al., 2019). Similarly, cells undergoing PAK1 or PAK2 gene knockout were viable, and their overall metabolic activity was usually comparable with that of the control cells (Fig. 6B), except for a few experiments with PAK2 KO. This finding would suggest that neither PAK1 nor PAK2 are specifically required for cell metabolism, either due to PAK1/PAK2 redundancy or thanks to the cell’s ability to overcome the lack of PAK activity by activating other signaling pathways on a longer time scale. However, detailed analysis of HeLa clones derived from one sample after PAK2 KO showed that although the efficiency of PAK2 gene targeting was high (11 of 12 clones had no or very weak signal from the PAK2 N-term antibody), a kinase-active modified PAK2 was detected in all of the clones by the phospho-specific PAK antibody (Supplementary Fig. S7A) as well as by an alternative PAK2 antibody targeting the C-terminal half. It thus seems that at least partly preserved PAK2 activity is necessary for long-term cell survival. This hypothesis is consistent with the fact that we were not able to obtain viable OCI-AML3 cells with PAK2 KO.

The majority of specific interaction points between PAK signaling and cell metabolic pathways reported to date were localized at the level of glucose uptake or glucose conversion to pyruvate (Fig. 1). In our model systems, PAK inhibition by IPA-3 did not consistently lower glucose uptake (Fig. 4A). Furthermore, IPA-3 significantly lowered the OCR even under glycolytic stress conditions, i.e., in the absence of glucose and pyruvate as fuels, both in adherent and leukemia cell lines (Fig. 4B). Thus, PAK targeting slows down cell metabolism regardless of glucose availability, and further research is needed to describe the mechanisms of PAK regulatory function(s) in the cell metabolic pathways. mTOR and PKM2, known metabolic regulators, are important mediators of PAK1 effects on glioma cell proliferation, migration, and drug resistance (Kim et al., 2020). Alternatively, inhibition of mTOR complex 1 with rapamycin or siRNA-mediated downregulation of raptor resulted in increased expression and activity of PAK1 in the lymphocytic Jurkat cell line (Rouquette-Jazdanian et al., 2015). Reduced expression of rictor, which forms part of mTOR complex 2, correlated with increased PAK1 activity in macrophages treated with fingolimod (Chen, W., Ghobrial, Li, & Kloc, 2018). In contrast, higher PAK1 activity resulting from mTOR activation was described in prostate cancer cells (Wang et al., 2017).

PAK group I proteins are key regulators of cytoskeletal dynamics. Emerging evidence points to a number of interconnections between cell adhesion and/or migration on the one hand and cell metabolic preferences on the other hand. A migratory phenotype of cancer cells, which is known to be promoted, for example, by c-Src-mediated PAK1 activation, is generally associated with increased glycolysis. In this sense, our finding that PAK1 inhibition by gene knockout increases the glycolytic rate (Fig. 6B,D) is counter-intuitive. However, the respective roles of the individual members of PAK group I, including different PAK1 splicing isoforms, are far from elucidated, even in adhesion signaling. We showed previously that both PAK2 and the dominant PAK1 isoform, PAK1Δ15, localize to focal adhesions and may thus participate in their assembly and/or turnover. In contrast, full-length PAK1 is involved in membrane protrusion formation (Parrini, Camonis, Matsuda, & de Gunzburg, 2009; Sells et al., 1997). The results presented in the current study suggest that PAK2 is a key player in the regulation of cell metabolism. Consistently, PAK2 is the most abundant PAK group I member in leukemia cells, which are very sensitive to PAK inhibition by IPA-3 (Kuželová et al., 2021).

## Conclusions

In conclusion, our results show that PAK group I is involved in the regulation of both mitochondrial respiration and aerobic glycolysis. Pharmacological inhibition of PAK group I blocks cell metabolism, and this effect is mainly mediated by inhibition of PAK nonkinase function(s). Although the roles of PAK1 and PAK2 are likely partly redundant, PAK2 activity is specifically associated with a shift of cell metabolic preferences toward glycolysis. This finding is consistent with a recent report indicating that PAK2 stimulates the activity of the cancer-specific isoform of pyruvate kinase, PKM2, which directs cell metabolism to aerobic glycolysis (Gupta et al., 2018). Alternatively, PAK1 activity was associated with glycolysis inhibition. Our results suggest the existence of important yet undescribed crosstalk between PAK signaling and metabolic pathways, situated within the Krebs cycle and/or within mitochondrial respiration. However, the individual roles of the different PAK group I members remain to be explored, both in adhesion signaling and in cell metabolic processes.

## Acknowledgment

The work was financially supported by the European Regional Development Fund, by the state budget of the Czech Republic (project AIIHHP: CZ.02.1.01/0.0/0.0/16_025/0007428, OP RDE, Ministry of Education, Youth and Sports), and by the Ministry of Health of the Czech Republic (project for conceptual development of the research organization No 00023736). The authors wish to thank M.Voráčová for expert technical assistance.

## Data availability statement

The data that support the findings of this study are available from the corresponding author upon reasonable request.

## References

Binder, P., Wang, S., Radu, M., Zin, M., Collins, L., Khan, S., … Liu, W. (2019). Pak2 as a Novel Therapeutic Target for Cardioprotective Endoplasmic Reticulum Stress Response. Circulation Research, 124(5), 696–711. 10.1161/CIRCRESAHA.118.312829 [doi]

Campbell, H. K., Salvi, A. M., O’Brien, T., Superfine, R., & DeMali, K. A. (2019). PAK2 links cell survival to mechanotransduction and metabolism. The Journal of Cell Biology, 218(6), 1958–1971. 10.1083/jcb.201807152 [doi]

Chen, W. L., Wang, J. H., Zhao, A. H., Xu, X., Wang, Y. H., Chen, T. L., … Jia, W. (2014). A distinct glucose metabolism signature of acute myeloid leukemia with prognostic value. Blood, 124(10), 1645–1654. 10.1182/blood-2014-02-554204 [doi]

Chen, W., Ghobrial, R. M., Li, X. C., & Kloc, M. (2018). Inhibition of RhoA and mTORC2/Rictor by Fingolimod (FTY720) induces p21-activated kinase 1, PAK-1 and amplifies podosomes in mouse peritoneal macrophages. Immunobiology, 223(11), 634-647. S0171-2985(18)30046-9 [pii]

Cheng, T. Y., Yang, Y. C., Wang, H. P., Tien, Y. W., Shun, C. T., Huang, H. Y., … Hua, K. T. (2018). Pyruvate kinase M2 promotes pancreatic ductal adenocarcinoma invasion and metastasis through phosphorylation and stabilization of PAK2 protein. Oncogene, 37(13), 1730–1742. 10.1038/s41388-017-0086-y [doi]

Coniglio, S. J., Zavarella, S., & Symons, M. H. (2008). Pak1 and Pak2 mediate tumor cell invasion through distinct signaling mechanisms. United States:10.1128/MCB.01532-07

Deacon, S. W., Beeser, A., Fukui, J. A., Rennefahrt, U. E., Myers, C., Chernoff, J., & Peterson, J. R. (2008). An isoform-selective, small-molecule inhibitor targets the autoregulatory mechanism of p21-activated kinase. England:10.1016/j.chembiol.2008.03.005

Grebenova, D., Holoubek, A., Roselova, P., Obr, A., Brodska, B., & Kuzelova, K. (2019). PAK1, PAK1Delta15, and PAK2: similarities, differences and mutual interactions. England:10.1038/s41598-019-53665-6 [doi]

Gupta, A., Ajith, A., Singh, S., Panday, R. K., Samaiya, A., & Shukla, S. (2018). PAK2-c-Myc-PKM2 axis plays an essential role in head and neck oncogenesis via regulating Warburg effect. England:10.1038/s41419-018-0887-0 [doi]

Gururaj, A., Barnes, C. J., Vadlamudi, R. K., & Kumar, R. (2004). Regulation of phosphoglucomutase 1 phosphorylation and activity by a signaling kinase. Oncogene, 23(49), 8118-8127.1207969 [pii]

Han, T., Kang, D., Ji, D., Wang, X., Zhan, W., Fu, M., … Wang, J. B. (2013). How does cancer cell metabolism affect tumor migration and invasion? Cell Adhesion & Migration, 7(5), 395–403. 10.4161/cam.26345 [doi]

Herst, P. M., Howman, R. A., Neeson, P. J., Berridge, M. V., & Ritchie, D. S. (2011). The level of glycolytic metabolism in acute myeloid leukemia blasts at diagnosis is prognostic for clinical outcome. Journal of Leukocyte Biology, 89(1), 51–55. 10.1189/jlb.0710417 [doi]

Jiang, Y., & Nakada, D. (2016). Cell intrinsic and extrinsic regulation of leukemia cell metabolism. International Journal of Hematology, 103(6), 607–616. 10.1007/s12185-016-1958-6 [doi]

Kanumuri, R., Saravanan, R., Pavithra, V., Sundaram, S., Rayala, S. K., & Venkatraman, G. (2020). Current trends and opportunities in targeting p21 activated kinase-1(PAK1) for therapeutic management of breast cancers. Gene, 760, 144991. S0378-1119(20)30660-0 [pii]

Kim, J. H., Seo, Y., Jo, M., Jeon, H., Kim, Y. S., Kim, E. J., … Suk, K. (2020). Interrogation of kinase genetic interactions provides a global view of PAK1-mediated signal transduction pathways. The Journal of Biological Chemistry, 295(50), 16906–16919. S0021-9258(17)50587-6 [pii]

Kreuzaler, P., Panina, Y., Segal, J., & Yuneva, M. (2020). Adapt and conquer: Metabolic flexibility in cancer growth, invasion and evasion. Molecular Metabolism, 33, 83–101. S2212-8778(19)30906-8 [pii]

Kumar, R., Sanawar, R., Li, X., & Li, F. (2017). Structure, biochemistry, and biology of PAK kinases. Netherlands: Elsevier B.V. S0378-1119(16)30981-7 [pii]

Kuzelova, K., Grebenova, D., Holoubek, A., Roselova, P., & Obr, A. (2014). Group I PAK inhibitor IPA-3 induces cell death and affects cell adhesivity to fibronectin in human hematopoietic cells. United States:10.1371/journal.pone.0092560 [doi]

Kuželová, K., Obr, A., Röselová, P., Grebeňová, D., Otevřelová, P., Brodská, B., & Holoubek, A. (2021). Group I p21-activated kinases in leukemia cell adhesion to fibronectin. Cell Adhesion & Migration, 15(1), 18–36. 10.1080/19336918.2021.1872760 [doi]

Licciulli, S., Maksimoska, J., Zhou, C., Troutman, S., Kota, S., Liu, Q., … Kissil, J. L. (2013). FRAX597, a small molecule inhibitor of the p21-activated kinases, inhibits tumorigenesis of neurofibromatosis type 2 (NF2)-associated Schwannomas. United States:10.1074/jbc.M113.510933 [doi]

Lu, J., Tan, M., & Cai, Q. (2015). The Warburg effect in tumor progression: mitochondrial oxidative metabolism as an anti-metastasis mechanism. Cancer Letters, 356(2 Pt A), 156–164. S0304-3835(14)00207-9 [pii]

Pandolfi, A., Stanley, R. F., Yu, Y., Bartholdy, B., Pendurti, G., Gritsman, K., … Steidl, U. (2015). PAK1 is a therapeutic target in acute myeloid leukemia and myelodysplastic syndrome. United States: by The American Society of Hematology.10.1182/blood-2014-12-618801 [doi]

Parrini, M. C., Camonis, J., Matsuda, M., & de Gunzburg, J. (2009). Dissecting activation of the PAK1 kinase at protrusions in living cells. United States:10.1074/jbc.M109.015271 [doi]

Rane, C. K., & Minden, A. (2019). P21 activated kinase signaling in cancer. Seminars in Cancer Biology, 54, 40–49. S1044-579X(17)30249-3 [pii]

Rouquette-Jazdanian, A. K., Kortum, R. L., Li, W., Merrill, R. K., Nguyen, P. H., Samelson, L. E., & Sommers, C. L. (2015). miR-155 Controls Lymphoproliferation in LAT Mutant Mice by Restraining T-Cell Apoptosis via SHIP-1/mTOR and PAK1/FOXO3/BIM Pathways. PloS One, 10(6), e0131823. 10.1371/journal.pone.0131823 [doi]

Sells, M. A., Knaus, U. G., Bagrodia, S., Ambrose, D. M., Bokoch, G. M., & Chernoff, J. (1997). Human p21-activated kinase (Pak1) regulates actin organization in mammalian cells. England:S0960-9822(97)70091-5 [pii]

Senapedis, W., Crochiere, M., Baloglu, E., & Landesman, Y. (2016). Therapeutic Potential of Targeting PAK Signaling. Netherlands:ACAMC-EPUB-68116 [pii]

Shalom-Barak, T., & Knaus, U. G. (2002). A p21-activated kinase-controlled metabolic switch up-regulates phagocyte NADPH oxidase. United States:10.1074/jbc.M206650200 [doi]

Sousa, B., Pereira, J., & Paredes, J. (2019). The Crosstalk Between Cell Adhesion and Cancer Metabolism. International Journal of Molecular Sciences, 20(8), 1933. doi: 10.3390/ijms20081933. 10.3390/ijms20081933 [doi]

Tsuji-Takayama, K., Kamiya, T., Nakamura, S., Matsuo, Y., Adachi, T., Tsubota, T., … Minowada, J. (1994). Establishment of multiple leukemia cell lines with diverse myeloid and/or megakaryoblastoid characteristics from a single Ph1 positive chronic myelogenous leukemia blood sample. JAPAN:

Tunduguru, R., Chiu, T. T., Ramalingam, L., Elmendorf, J. S., Klip, A., & Thurmond, D. C. (2014). Signaling of the p21-activated kinase (PAK1) coordinates insulin-stimulated actin remodeling and glucose uptake in skeletal muscle cells. Biochemical Pharmacology, 92(2), 380-388. https://doi.org/10.1016/j.bcp.2014.08.033

Varshney, P., & Dey, C. S. (2016). P21-activated kinase 2 (PAK2) regulates glucose uptake and insulin sensitivity in neuronal cells. Molecular and Cellular Endocrinology, 429, 50–61. S0303-7207(16)30092-2 [pii]

Wang, Z., Jia, G., Li, Y., Liu, J., Luo, J., Zhang, J., … Chen, G. (2017). Clinicopathological signature of p21-activated kinase 1 in prostate cancer and its regulation of proliferation and autophagy via the mTOR signaling pathway. Oncotarget, 8(14), 22563–22580. 10.18632/oncotarget.15124 [doi]

