## Supplementary Figures for "PAK1 and PAK2 in cell metabolism regulation"

### Supplementary Figures S1 to S9

Figure S1: Example of metabolic measurement in HEK293T cells. Cells (20k/well) were seeded into a Seahorse plate and incubated for 24 h. After medium change, they were treated with 20 µM IPA-3 (closed circles) or with 20 µM PIR3.5 (triangles) in duplicate for 1 h. The wells were washed twice in Sehorse medium, and the metabolic rates were measured using the Seahorse XFp device. The points represent the means and standard deviations of duplicate wells. Injections of oligomycin (OM), FCCP and blocking compounds are indicated by black arrows. The background values were taken from the last recorded points. The basal OCR and ECAR levels were calculated from the values just before OM addition, the maximal values were obtained from the maximal values reached after OM addition, as it is illustrated for the control sample (open circles).

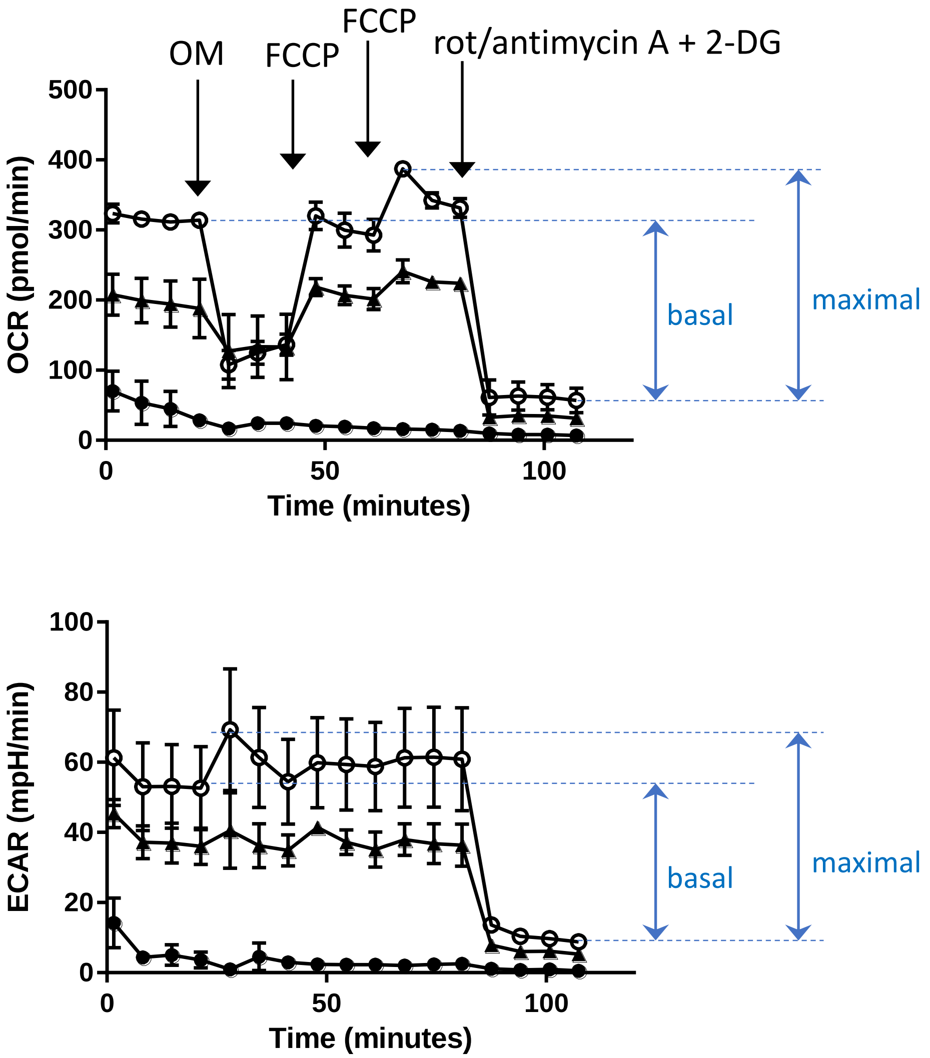

Figure S2: Effect of 5 h FRAX597 treatment on OCI-AML3 cell metabolism. The cells were incubated for 4 h with FRAX597 at 2 µM or 10 µM concentration, then seeded on CellTak in Seahorse medium. FRAX597 was added, and the cells were incubated for an additional 1 h prior to the analysis.

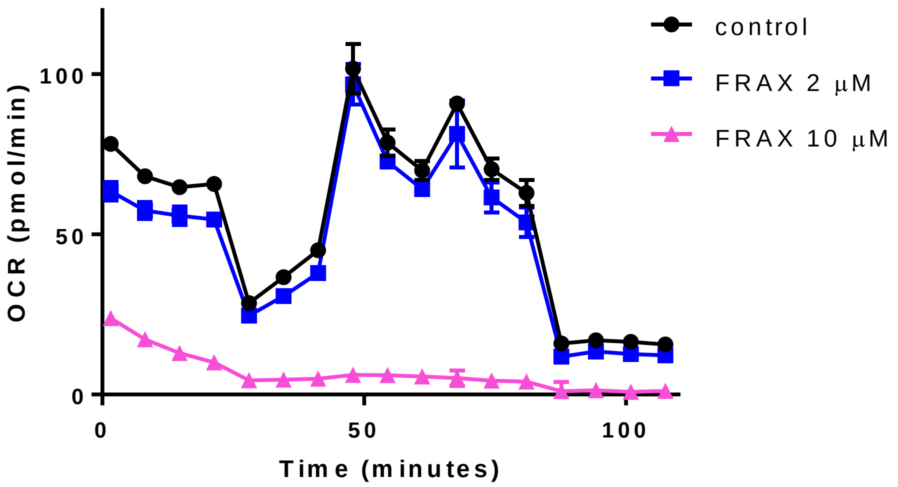


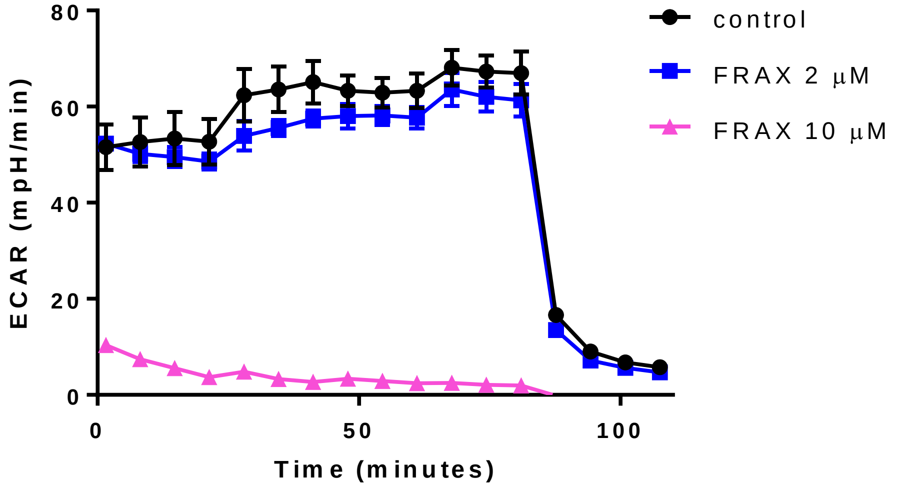

Figure S3-panel A: Results of experiments with primary AML cells.
The cells were pretreated with 20 µM IPA-3 or PIR3.5 for 1-2h at 3x10$^5$/ml cell density, washed, and seeded into Seahorse plate at 4x10$^6$ /ml (i.e., 2x10$^5$ cells/well). The points show means and s.d. of well duplicates. Results from freshly isolated primary cells of 4 different patients with AML at diagnosis. Sequential injections of oligomycin (OM), FCCP 0.3 and 0.5 µM, and rotenone + antimycin A + 2-deoxyglucose (rot/A + 2-DG) are indicated by the arrows. The protocol was the same in all cases, except for missing 2-DG in the first one.

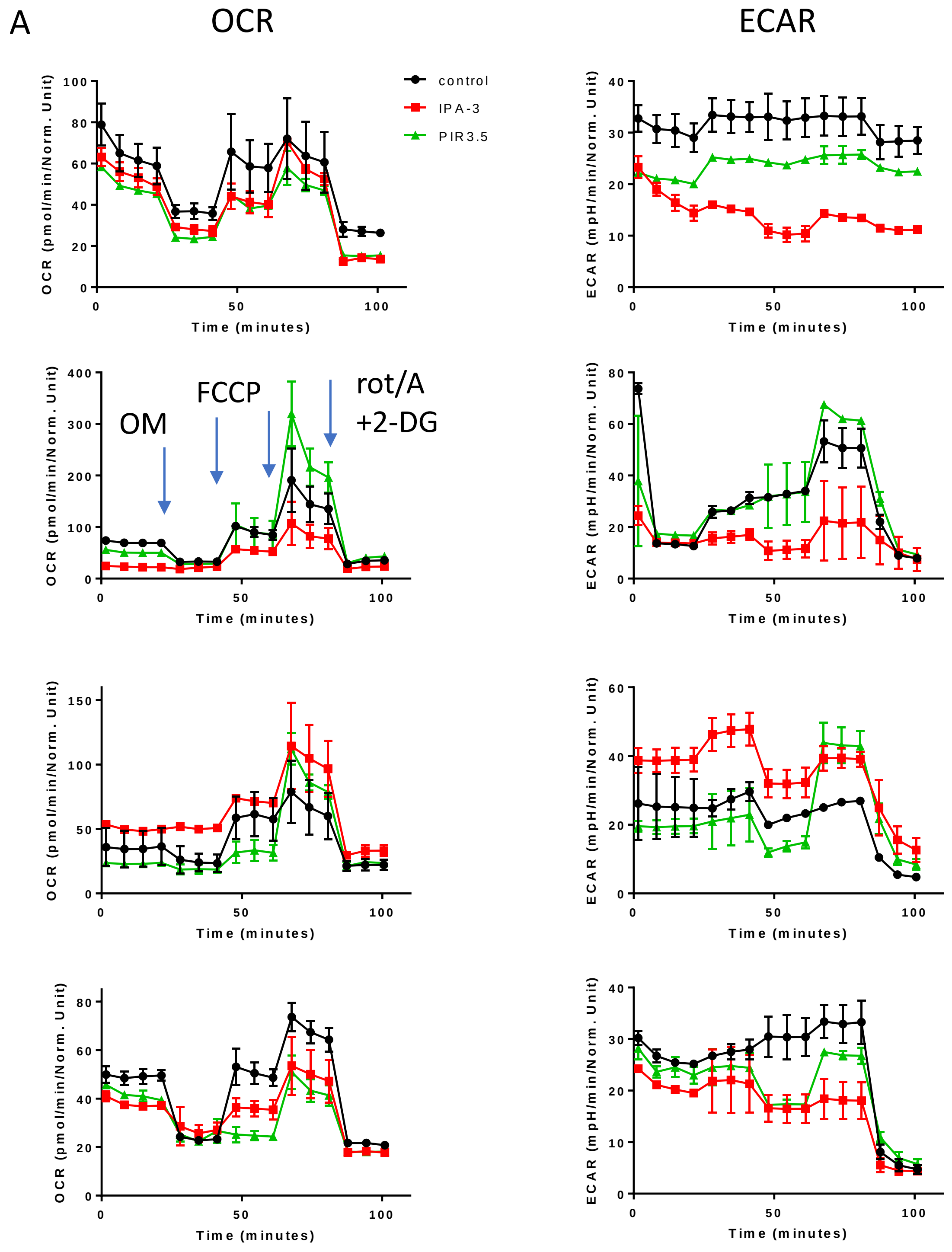

Figure S3-panel B: Results of experiments with primary AML cells.
The cells were treated with 50 µM IPA-3 or PIR3.5, either in the same settings as in Fig. S3-A or during incubation in Seahorse plate as in Fig. 3 for cell lines. Results obtained with samples from two different patients with AML at diagnosis. The protocol for Seahorse XFp was the same as in Fig. S3-A.

B

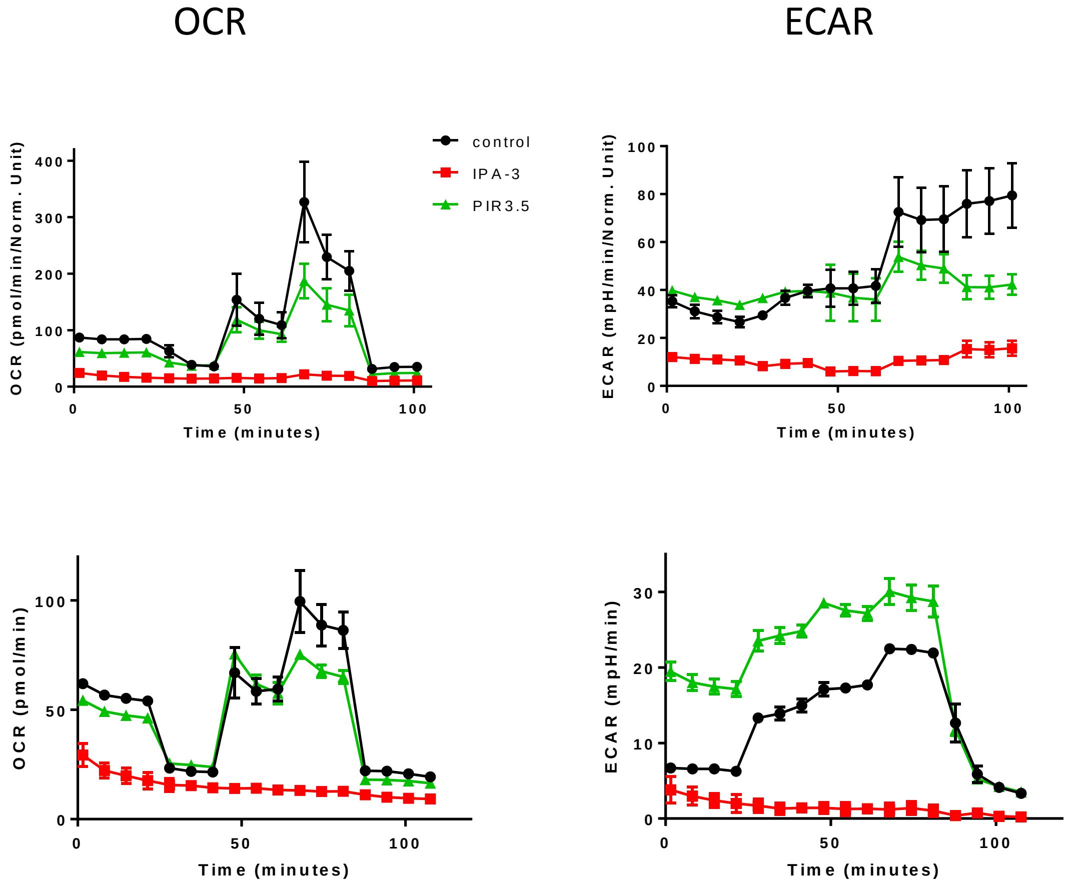

Figure S4: Examples of Seahorse XFp records for cells under glycolytic stress. The cells were seeded in glucose-free and pyruvate-free medium and incubated for 1 h in the presence of 20 µM IPA-3 or PIR3.5 prior to Seahorse XFp measurement. The graphs show means and s.d. from well duplicates. A: HEK293T cells, B: HeLa cells, C: Karpas-299 cells. Controls: black circles, IPA-3: red squares, PIR3.5: green triangles. Injections of glucose, oligomycin (OM), FCCP, rotenone/antimycin A (rot/A), and 2-deoxyglucose (2-DG) are indicated by the arrows.

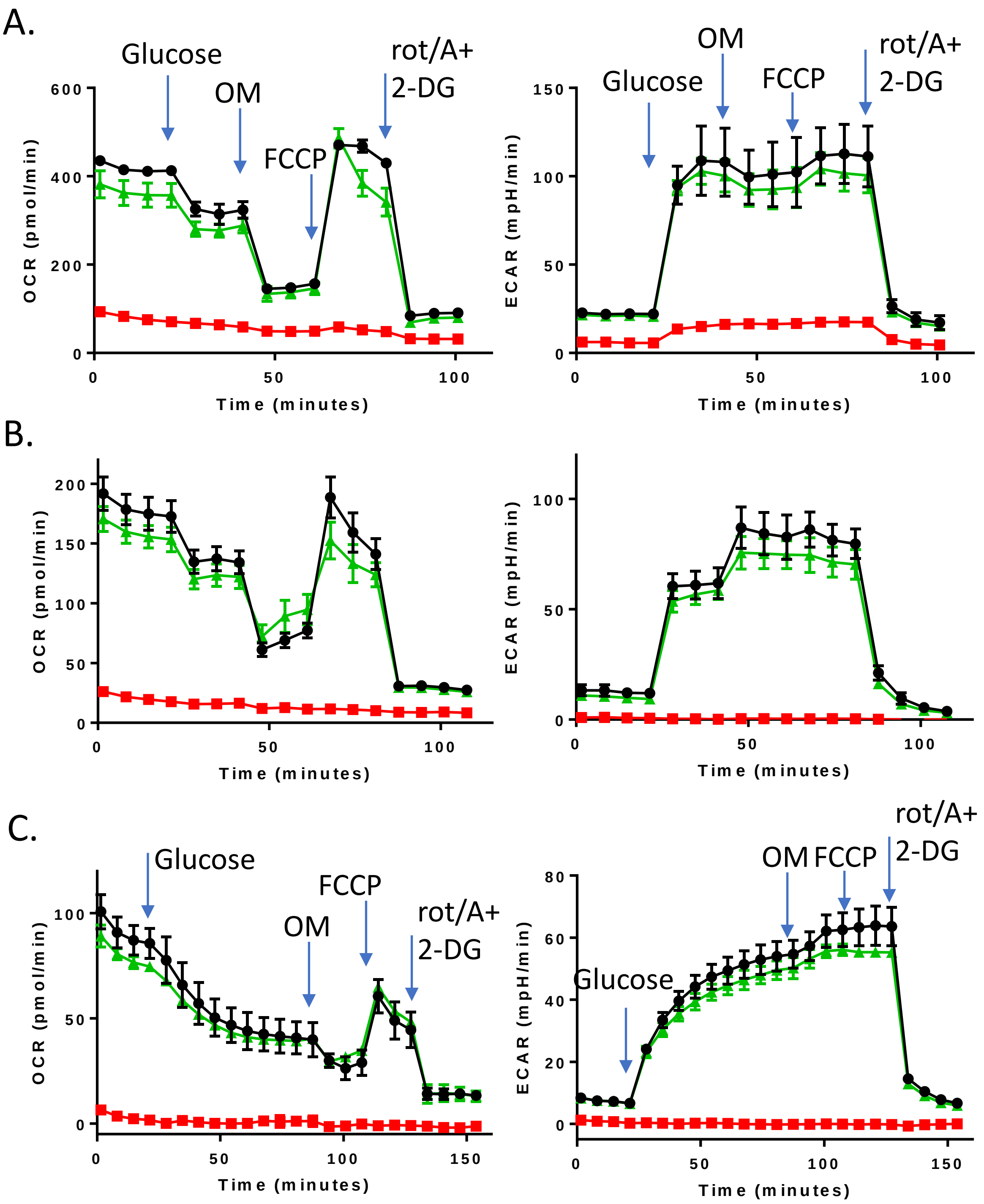

Figure S5: Correlation of siRNA PAK2 effects on PAK2 and PAK1 levels in individual experiments. HEK293T cells were transfected with siRNA PAK2 and the levels of PAK1, PAK2 and pSer144/141 PAK1/PAK2 were analyzed by western-blot. A: examples of western-blot results. B: The graphs show intensities of individual PAK1 bands (top) and pSer144 PAK1 bands (bottom), as a function of PAK2 levels. All values are expressed relative to those from control samples transfected with nontargeting (NT) siRNA. Equal loading was checked by β-actin staining.

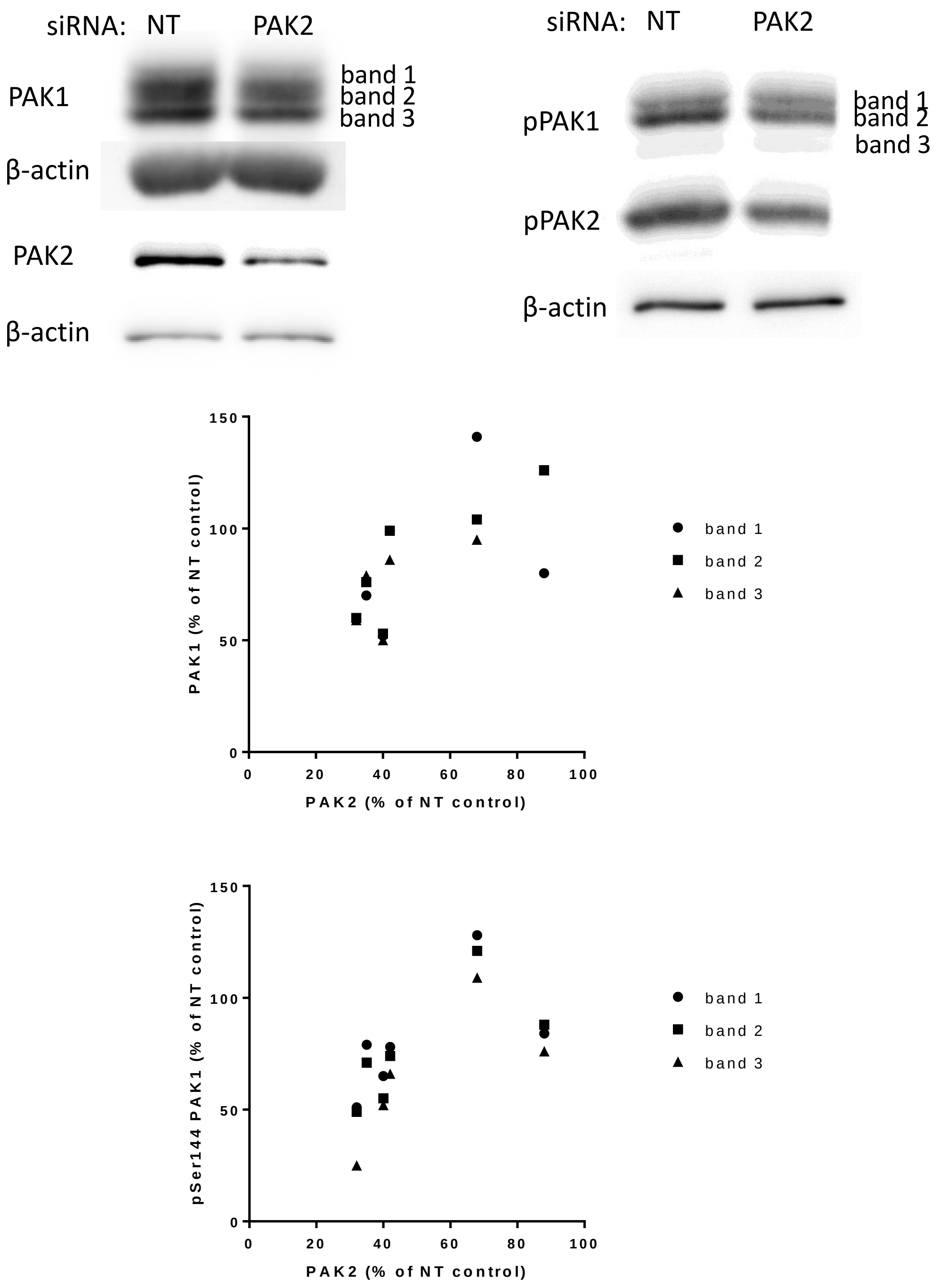

Figure S6: Effect of PAK targeting by siRNA on the cell respiration rate under glycolytic stress.

The cells were transfected with nontargeting (NT) siRNA or PAK1/PAK2 siRNA and cultured for 24 h under standard conditions. Then, they were harvested, and aliquots were seeded into microtitration plates in triplicate (NT siRNA and PAK1/PAK2 siRNA) and cultured for further 24 h. The cells were washed in glucose/pyruvate-free medium and incubated for 1 h prior to Seahorse measurement. The cell metabolic rates are given as relative to the corresponding NT siRNA control and represent means and standard deviations of 3 to 4 independent experiments for each siRNA and each cell line.

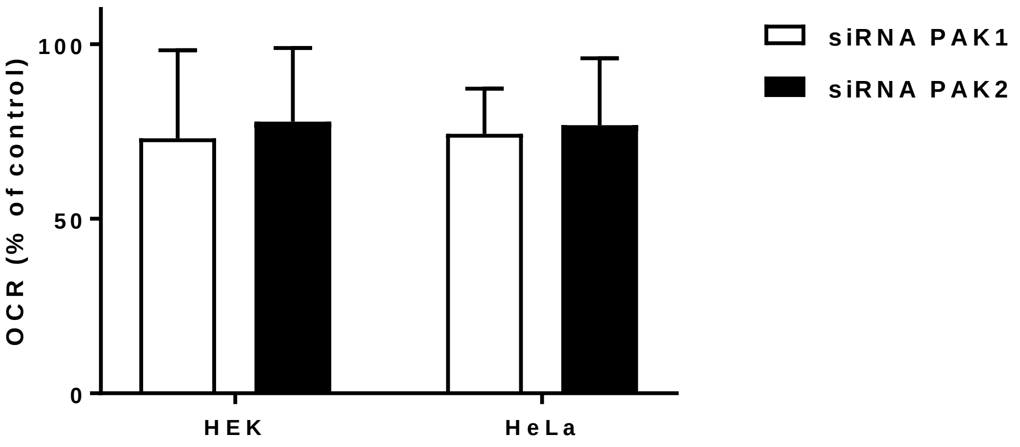

<u>Figure S7</u>: Properties of HeLa clones with PAK2 knockout.
A. PAK expression in individual clones established from HeLa cells after CRISPR/Cas9-induced PAK2 modification. B. Clone D3 was transfected with PAK2-GFP plasmid and the signal from GFP was analyzed by flow cytometry in the presence of propidium iodide (PI) to exclude dead cells. C. The metabolic rates of the clone D12 transfected or not with PAK2-GFP (means and s.d. from 4 experiments).

A. HeLa clones grown from PAK2 KO

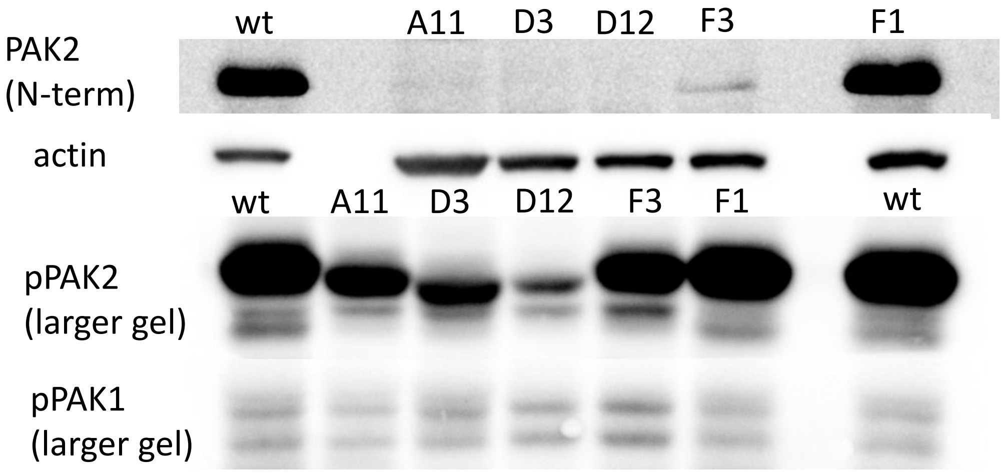


B. HeLa clone D3

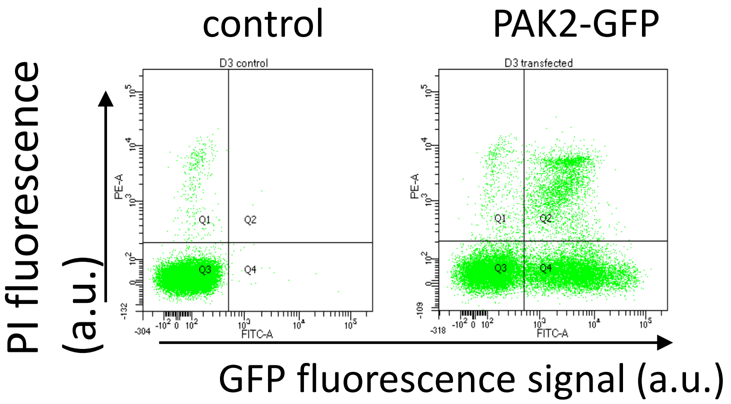


C. HeLa clone D12 transfected with PAK2-GFP plasmid

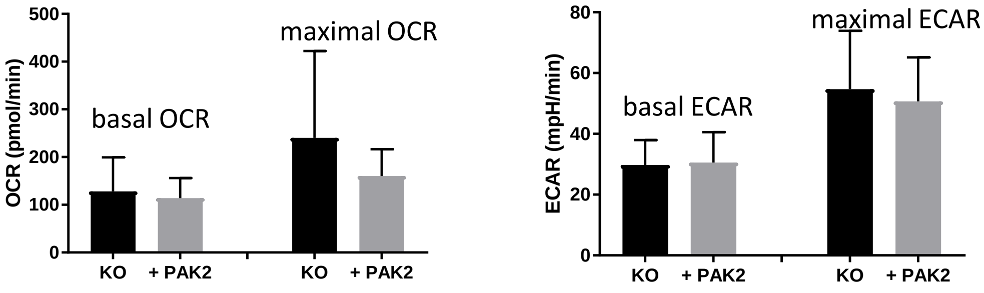

Figure S8: Effect of Src kinase inhibition by dasatinib on the cell metabolic rates.
HeLa or HEK293T cells were treated with 100 nM dasatinib for 1 h prior to analysis by Seahorse XFp device. The basal OCR and ECAR values from treated samples were expressed relative to the corresponding values from untreated controls. The bars show means and s.d. from 4 independent experiments for HeLa, resp. 3 for HEK293T cells, each performed in triplicate.

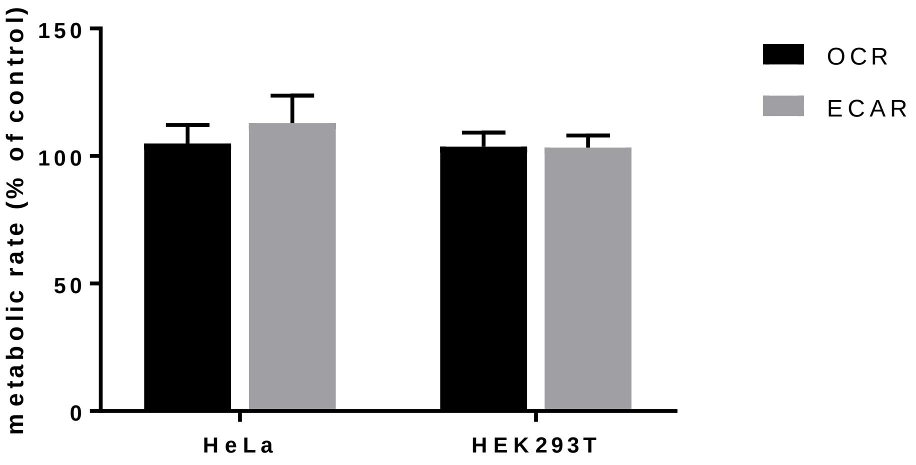

Figure S9: Effect of FRAX597+PIR3.5 combination on cell metabolic rates. HeLa, HEK293T, or OCI-AML3 cells were treated with 10 µM FRAX, 20 µM PIR3.5 or with the combination for 1 h prior to analysis by Seahorse XFp device. The points show means and s.d. from well replicates. Injections of oligomycin (OM, 1 µM), FCCP (0.3 and 0.5 µM final), rotenone/antimycin A (rot/A, 0.5 µM), and 2-deoxyglucose (2-DG) are indicated by the arrows.

### A. HEK293T

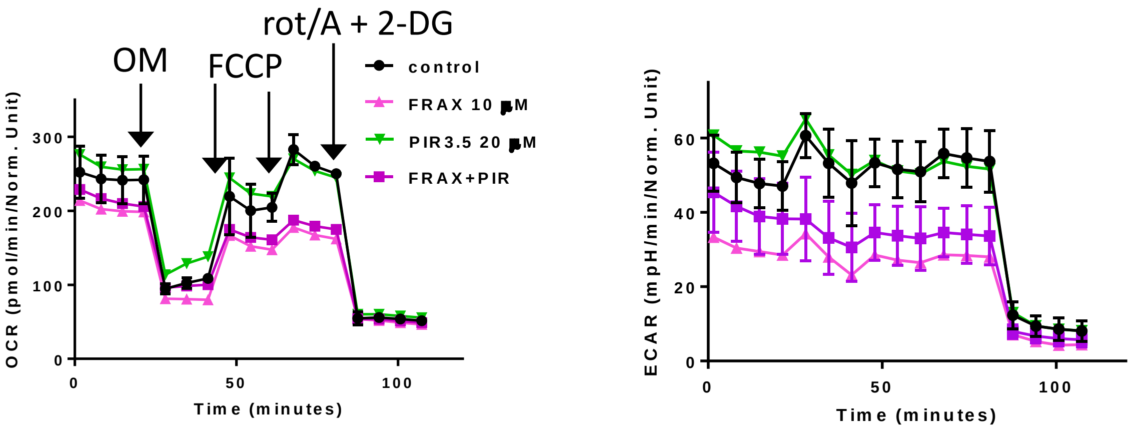


### B. HeLa

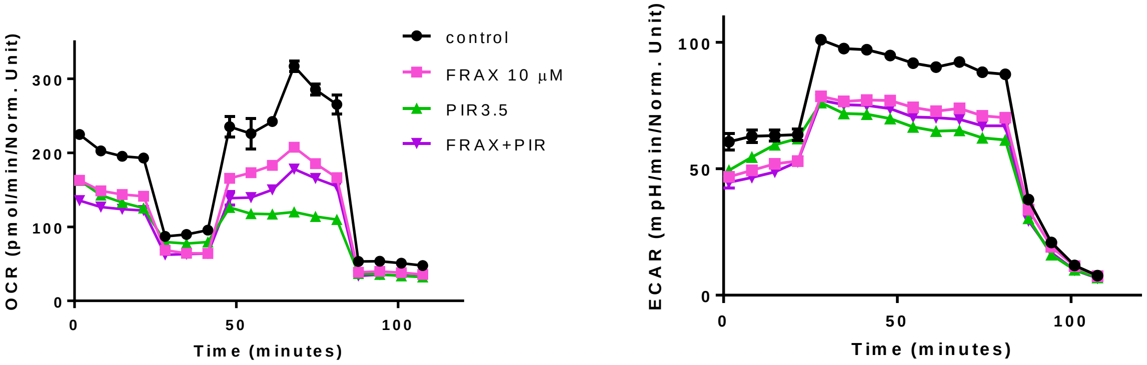


### C. Karpas-299

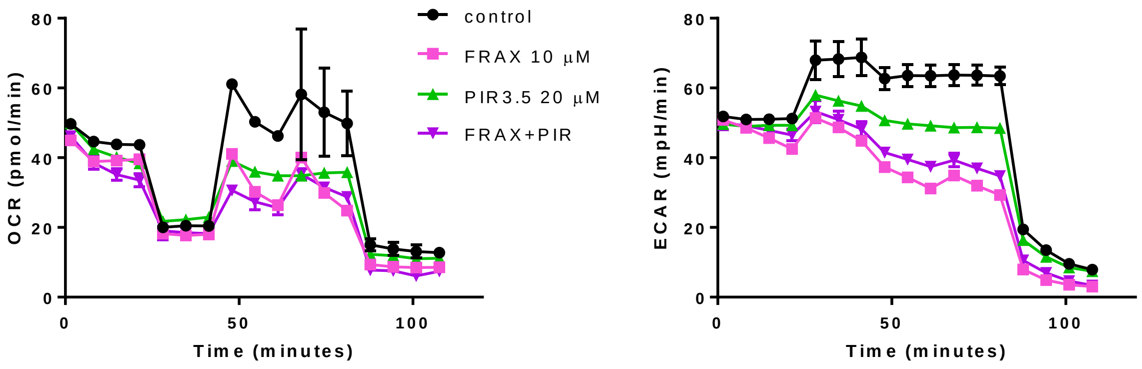
